# A Charge-Encoded Rheostat Permits Helix Nucleation but Limits Propagation in Skp1

**DOI:** 10.64898/2026.07.13.737693

**Authors:** Debarghya Mitra, Simran Tolani, Amrita Bhattacharya, Supriya Prathihar, Sagarjyoti Pathak, Arpita Prasad, Christian Griesinger, Ashutosh Kumar, Sarath Chandra Dantu

## Abstract

Molecular recognition by intrinsically disordered regions (IDRs) is widely thought to involve coupled folding and binding, yet the sequence features that regulate this transition remain underexplored. Here we show that helix 8 (H8), a disordered C-terminal segment of the SCF ubiquitin ligase adaptor Skp1, is intrinsically prevented from forming a stable helix by its own sequence grammar. Using an integrative approach to dissect its conformational dynamics, we find that H8 frequently nucleates helical structure but rarely propagates into a fully formed stable helix, populating instead a shallow metastable basin of helical intermediates. Contrary to conventional models of helix-coil exchange, where nucleation is rate limiting, helix initiation in H8 is readily accessible, while propagation is selectively suppressed by a glutamate-rich acidic patch. This acidic segment acts as a charge-sensitive conformational rheostat that limits helix extension and maintains H8 in a predominantly disordered state. As a result, H8 transiently samples partially helical conformations on the microsecond timescale without committing to a stable fold. We propose that this propagation-limited mechanism preserves conformational flexibility while maintaining recognition competence across a structurally diverse family of F-box binding partners. More broadly, our findings suggest that charged-hydrophobic-charged sequence patterning can encode conditional, context-dependent structure as a general organisational principle in intrinsically disordered proteomes.

## Introduction

Proteins that function as hubs in cellular networks frequently lack a single, well-defined tertiary structure^1,2^. Their functional versatility arises from the ability to interconvert among multiple conformations, consistent with the paradigm of single sequence-many structures^3–5^. Such proteins or protein segments that lack stable tertiary organization are classified as intrinsically disordered proteins (IDPs) or intrinsically disordered regions (IDRs), respectively and constitute nearly ∼60% of the eukaryotic proteome^6,7^. It is well known that mutations in the disordered regions of IDPs/IDRs can lead to pathogenic conditions like Alzheimer’s, Parkinsons, and even,cancer^8,9^.

Unlike structured proteins, IDPs traverse a continuum of states between the classical classes of secondary structures (helix, beta-sheet and coil), underscoring the need for approaches that resolve dynamic ensembles rather than time-averaged conformations^10^. AI-based structure prediction tools such as AlphaFold2^11^ and others^12,13^ have revolutionized structure determination for globular proteins, yet they fall short on predicting structures of metamorphic proteins^14,15^ and IDP/IDR^16^. Experimental methods, for structure elucidation of IDPs/IDRs have largely relied on static representations, with X-ray crystallography^17^ and cryo-EM^18^ predominantly capturing the stabilized conformation i.e. in complex with the partner protein. Solution state NMR captures the native disordered state of proteins^19,20^, but because of conformational exchange that spans multiple time scales^20^, experimental complexity poses a challenge for characterization of the largely invisible key intermediates required to study the underlying mechanism of structural transitions necessary for biological function^21–24^.

Among the plausible secondary structure transitions exhibited by IDRs/IDPs, helix-coil transitions are very prominent in many proteins like alpha-synuclein^25^, p27^Kip1^ ^26^,ACTR^27^, N-terminal domain (DBD) of the bacterial cytidine repressor (CytR)^28^, C-terminal tail of the measles virus nucleoprotein (NTAIL)^29^ to name a few. The classical Helix-Coil theory^30,31^, provide a thermodynamic framework to understand helix-coil transitions. It suggests that in helix formation, nucleation is the most demanding step, followed by easy helix propagation. However, these models largely treat helices as isolated structural elements as they only investigate short peptides^32^ and do not explicitly address the kinetics of proteins, their residue-level heterogeneity, or entropic contributions that dominate disorder-order transitions within IDRs. Furthermore, most of the studies have reported on the kinetics and mechanism of helix to coil transition but details on the reverse transition are missing. CD based studies, on short segments have demonstrated that environment^33,34^, is a key nucleating trigger for helix formation, we still lack atomistic level structural details of the process. Computational studies have quantified statistical weights of each residue and their position to estimate a thermodynamic model to determine nucleation energy ^35^. To summarise, we still lack explanation of how helices nucleate, unravel, and dynamically interconvert highlighting unresolved aspects of how helix-coil transitions are realised in intrinsically disordered systems.

For studying disorder–order coil-helix transitions within a functional IDR, S-phase kinase-associated protein-1 (Skp1) is an ideal model system (Figure 1A-B)^36–38^. It is an essential adaptor in the Skp1-Cullin1-F-box (SCF) E3 ligase multiprotein complex^39–41^ (Supplementary Figure 1), which mediates polyubiquitination of proteins meant for protein degradation^40^. Skp1 binds to ∼69 distinct F-box proteins (FBPs) ^42^and thereby participates in diverse cellular pathways ranging from cell-cycle control to inflammation^43^ and cancer^43,44^. The C-terminal helices helix6, helix7, and helix8 (H6-H8) from the C-terminal tail of Skp1 form the complex with F-box domain of the FBPs^39^. H8 of Skp1 has exhibited two distinct conformations. In the F-box bound Skp1 from X-ray crystallographic studies, H8 is a 10-residue helix^39^, whereas in the apo form of Skp1 C-terminal region becomes disordered^36^. Despite clear evidence for conformational heterogeneity and predicted coupled folding dynamics^37^, the kinetic mechanisms and intermediates that enable Skp1 to shift between helix or coil states is not yet understood.

**Figure 1:**
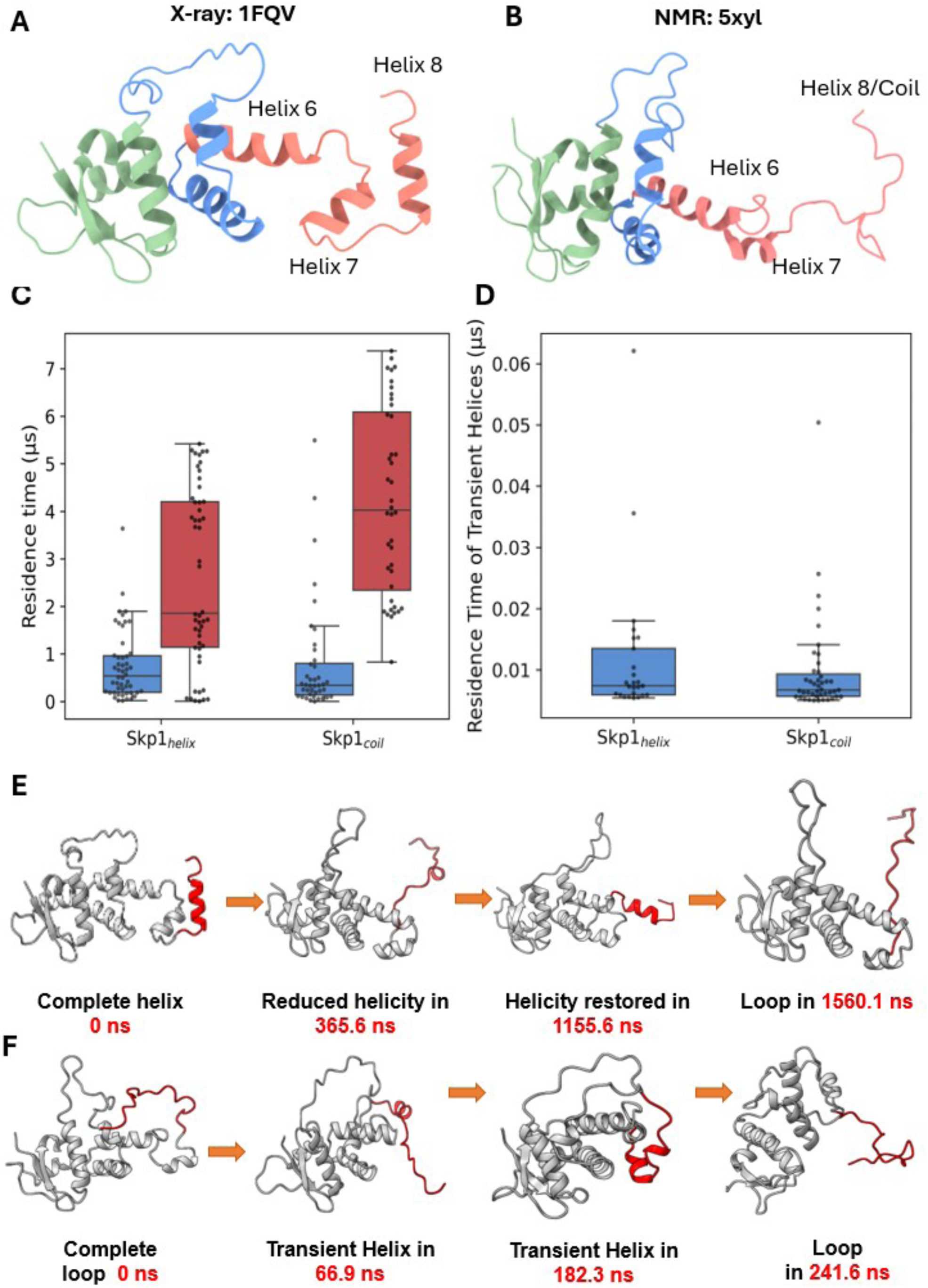
Starting conformations of Skp1 simulation trajectories, (A) Skp1_helix_ conformation with a fully formed helix (H8) at the C-terminal end, from PDB ID:1FQV and (B) Skp1_coil_ conformation with a disordered C-terminal end at, adapted from PDB ID:5XYL. (C)The proportion of time (residence time) residues in Helix 8 spent as helix(blue) or coil(red) distributed across Skp1_helix_ and Skp1_coil_ ensembles whereas the same residues when they form transient helices across the two ensembles are visualized through (D) suggesting a similar proportion sampled in both ensembles. Extraction of frames from a representative trajectory in Skp1_helix_ that follows the initial folding of H8 followed by a coil state and subsequent transition into an intermediate helix and finally culminating into a coil at different instances in the trajectory(E). A similar pathway extracted from a trajectory of Skp1_coil_ ensemble describes the formation of transient helices from coil before finally ending as a coil(F). In both Figures E-F timestamps of the frame, from which the conformation is extracted is included, but these formations are not happening at regular intervals of the trajectory.

In this work, we address these questions by combining long-timescale molecular dynamics (MD) simulations, Markov State Modelling (MSM), relaxation dispersion NMR and CD spectroscopy to resolve the helix-coil transition of the Skp1 C-terminal IDR at atomic resolution. We show that H8 does not undergo a simple two-state transition but instead populates a coil-dominated ensemble containing low-population, partially helical intermediates. These intermediates arise through frequent local nucleation but rarely propagate into a persistent full-length helix. We further show that the charged-hydrophobic-charged sequence architecture of H8 functions as a conformational rheostat that regulates this propagation-limited behaviour, maintaining a dynamically exchangeable ensemble capable of supporting F-box protein recognition without locking the region into a stable folded state.

## Results

An effective understanding of Skp1’s ability to engage multiple partners requires a detailed characterisation of its conformational dynamics. To investigate the mechanism of helix-coil interchange within Skp1’s C-terminal IDR, we performed MD simulations of apo-Skp1 initiated from two distinct structural states i.e., helix (Skp1_helix_; 50 replicas; 170µs) and coil (Skp1_coil_; 40 replicas; 190µs) using structures from X-ray (1FQV) ^39^and solution state NMR (5XYL)^36^ respectively (Supplementary Tables 2 and 3). To quantify kinetics and energetics of helix-coil interchange, we used MSM and validated the resulting timescales using extreme- CPMG^45^ (E-CPMG) relaxation dispersion experiments. Using circular dichroism (CD) on a synthetic H8 peptide (H8^syn^) we were able to probe the influence of sequence features, and how charged residues a regulate helix-coil interconversion. Consistent with previous studies^36,37^, Skp1 can be partitioned into a structurally stable core (helices H1-H5) and a flexible C-terminal region comprising H6-H8 (Supplementary Fig. 1, Table 1). Analysis of residue-level fluctuations confirms that conformational dynamics are largely confined to this C-terminal region (Supplementary Fig. 2), motivating a focused investigation of helix-coil transitions within H8.

### H8 exhibits asymmetric helix–coil kinetics with a dominant coil population

Simulation of both Skp1_helix_ and Skp1_coil_ trajectory ensembles reveal frequent helix-coil interconversion in H8 (residues 147-156: ‘EEEEAQVRKE’) based on secondary structure analysis. Independent of the starting conformation, both ensembles converge to a shared dynamic behaviour characterised by repeated transitions between helix and coil states (Fig. 1C, Supplementary Fig. 3). The coil state dominates the conformational ensemble, comprising 77.6% and 83.9% of the Skp1_helix_ and Skp1_coil_ trajectories respectively. In the Skp1_coil_ ensemble, 75% of trajectories remain predominantly coiled, with helix formation observed only intermittently, while in the Skp1_helix_ ensemble, 90% of trajectories undergo unfolding followed by helix-coil transitions (Supplementary Fig. 4).

Transition analysis highlights rapid interconversion between states, with both ensembles exhibiting multiple short-lived events (Skp1_helix_: 25 and Skp1_coil_: 43; median <0.01 μs; Fig. 1D). Despite the dominance of the coil state, the cumulative time required to form helices is comparable across both ensembles (∼2 μs), indicating frequent but short-lived helix formation events. Notably, helix-to-coil transitions occur readily, whereas coil-to-helix transitions are less frequent, indicating intrinsic kinetic asymmetry in the exchange process. Snapshots from representative trajectories further illustrate the rapid sub-microsecond interconversion between helix and coil states (Fig. 1E-F). H8 therefore populates a predominantly coil-like ensemble characterised by repeated transient helix formation and inefficient propagation into stable helices.

### Nucleation without propagation limits helix formation in H8

Helix-coil transitions within H8 predominantly generate short helical segments spanning 5-8 residues, with helices of length 5 and 7 representing the most populated states across both simulation ensembles (Fig. 2A-B). In the Skp1_helix_ ensemble, helices of length 5 and 7 comprise 53.1% and 20.6% of the observed helical states, respectively, whereas the Skp1_coil_ ensemble is strongly dominated by helices of length 5 (92.1%). These transient helices are short-lived, with most events exhibiting residence times below 0.2 μs. Although rare long-lived events are observed in the Skp1_coil_ ensemble (∼2.5 μs), fully propagated helices spanning the entire H8 segment are absent across both ensembles. The longest observed helices correspond to rare 8 residue states (<0.2%), indicating that helix formation remains restricted to partially propagated intermediates. Representative conformations extracted from the trajectories reveal progressive increases in intra-helical hydrogen bonding with increasing helix length (Fig. 2A- B). However, all transient helices ultimately revert to coil conformations, suggesting that local stabilisation alone is insufficient to maintain a persistent helical state. Helices of length seven and eight are seen in F-box bound Skp1 structures (Supplementary Figure 5).

**Figure 2:**
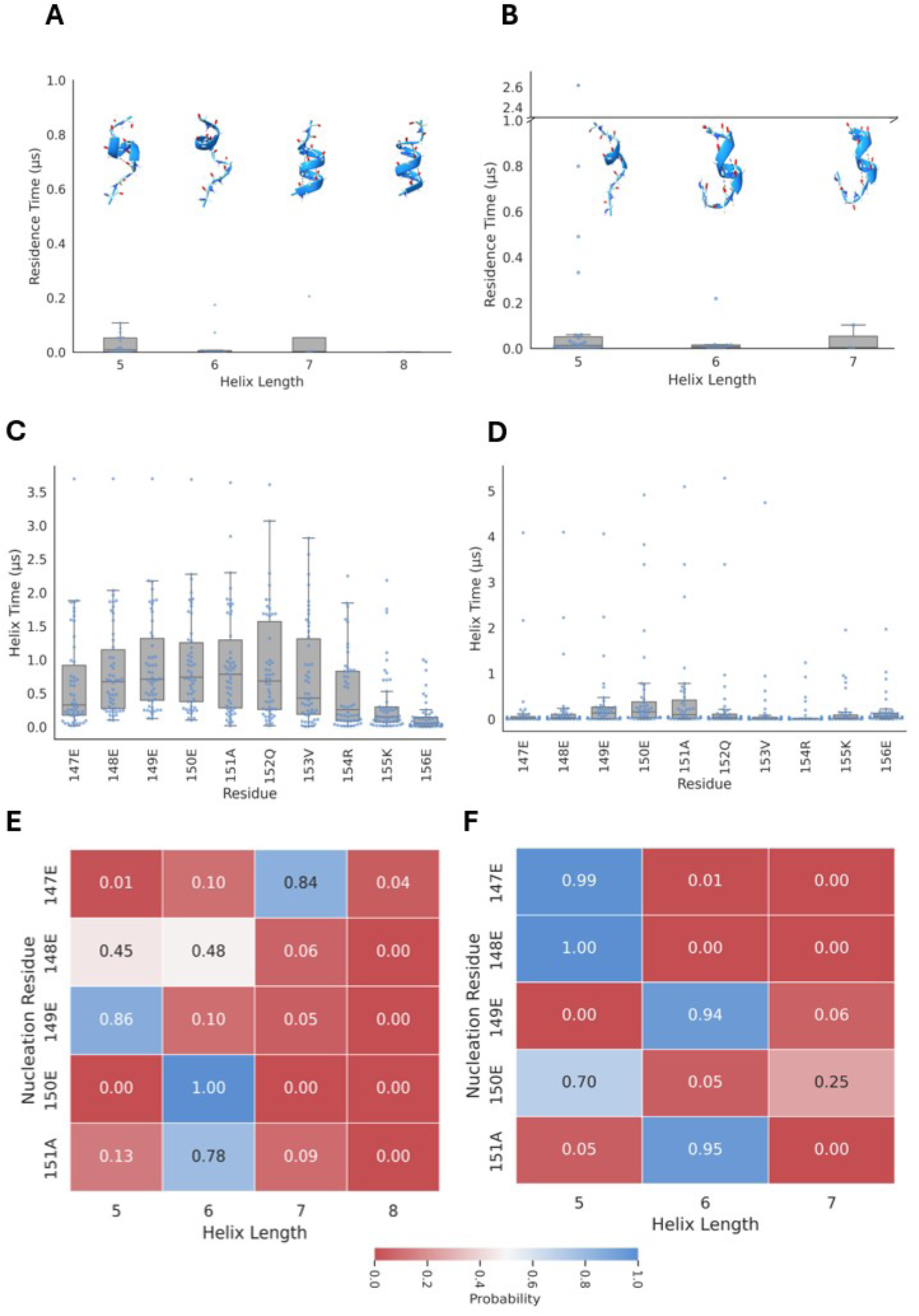
Distribution of variable lengths of transient helices from the two trajectory ensembles of Skp1_helix_ (A) and Skp1_coil_(B) highlights the time spent across each trajectory for a specific helix length. The occupancy time of each residue of H8 spent as a helix from the two trajectory ensembles of Skp1_helix_ (C) and Skp1_coil_ (D) from DSSP data highlights how each residue has a non-uniform trend and suggests a high occurrence of non-contiguous helix formation. The probability of formation of helices of variable lengths based on the residue that first commits to a transition from coil to helix across Skp1_helix_ (E) and Skp1_coil_ (F) trajectory ensembles (nucleation residue) suggests formation of helices from various starting points, confirming our context of “easy nucleation” and “limited propagation”

Residue-level secondary structure analysis further reveals heterogeneous helix propensity across H8 (Fig. 2C-D). H8 residues E149, E150, A151 and Q152 spend a large proportion of time (∼1 µs) as a helix (Supplementary Table 6). These residues exhibit the highest cumulative residence times within helical conformations, suggesting that local helix initiation is distributed across multiple neighbouring residues rather than arising from a single residue dominant nucleation site. The disparity in the distribution of helix time across the H8 residues (Figs 2C- D) suggests a non-cooperative nature of helix formation with many nucleation residues, but an inability to extend and form a stable helix.

This behaviour becomes more apparent when helix propagation is analysed as a function of the initiating residue (Fig. 2E-F). In both ensembles, helix initiation occurs across multiple positions within H8, but distinct residues exhibit reproducible preferences for specific helix lengths. For example, E147 preferentially gives rise to longer helices in the Skp1_helix_ ensemble, whereas E149 predominantly stabilises shorter helices. In contrast, the Skp1_coil_ ensemble exhibits more restricted propagation profiles, with most initiation events producing helices of length 5 or 6. Notably, the initiating residue does not strongly predict the final helix length, suggesting limited cooperativity between local nucleation and subsequent propagation. Together, these results support a model in which H8 undergoes frequent local helix nucleation but inefficient propagation into stable extended helices. Rather than forming a concerted all-or-none transition, helix formation proceeds through a heterogeneous ensemble of locally stabilised intermediate states characteristic of weakly cooperative folding within an IDR.

### Intermediate states mediate asymmetric helix-coil interchange in H8

Markov state model^46,47^ was carried out using writhe^48^, including just H8 residues (147-156) across both trajectory ensembles. MSM of the combined ensemble resolves the conformational landscape into six macrostates organised into two dominant basins connected by a set of intermediate states (Fig.3; Supplementary Fig. 6). Two macrostates (M0, 44.3%; M4, 36.7%) constitute the dominant coil basin, collectively accounting for ∼81% of the equilibrium population and consistent with the predominantly disordered ensemble observed by DSSP analysis (Fig. 3A). Both states are characterized by extended conformations lacking persistent secondary structure. In contrast, a single sparsely populated macrostate (M3, ∼0.1%) represents the fully propagated helical conformation corresponding to the starting Skp1_helix_ structure. Three intermediate macrostates (M2, 16.9%; M5, 1.8%; M1, 0.2%) connect the coil and helical basins and define the transition region of the landscape.

**Figure 3:**
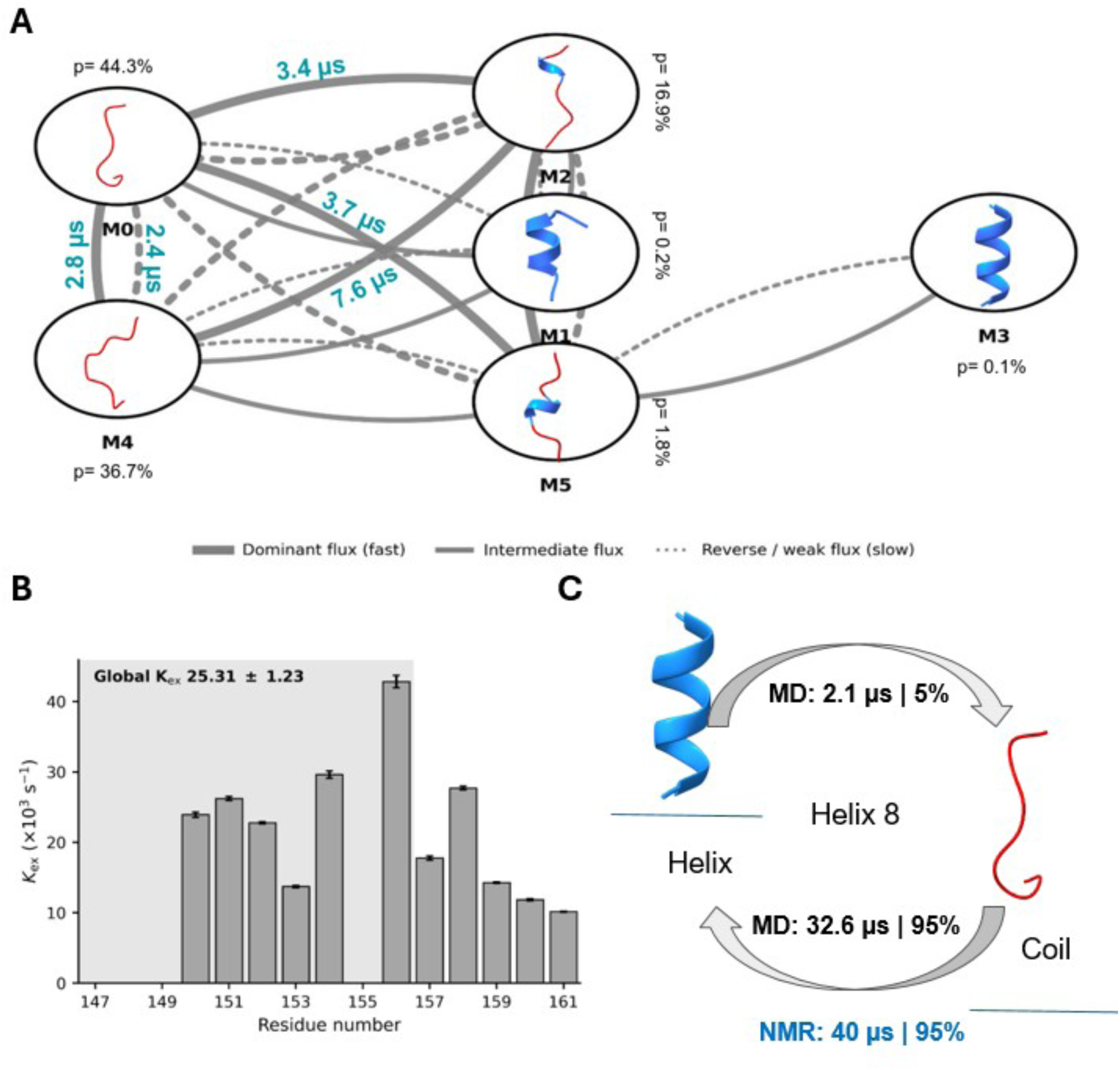
Conformational landscape of H8 was resolved into six major macrostates based on helicity (M0-M5) based on decomposing the length of H8 (147-156) from Skp1helix and Skp1coil trajectories(A). Solid arrow widths and dotted arrow widths between the macrostates suggest high/intermediate magnitude flux rates (based on width thickness) and reverse/weak flux rates (based on width thickness), respectively. Conformations have been extracted for each macrostate to highlight the diversity in helical content (in blue) or even coil (in red) for H8. Percent populations of each macrostate marked across each, indicate the predominance of coil conformations. Mean transition times are marked (in cyan) across the macrostates to highlight fast exchanging transitions, whereas slow transition times have been omitted from visual representation for clarity. Kinetic exchange rates extracted from E-CPMG experiment at 272K (B) include residues at the C-terminal end of Skp1. The shaded region in (B) marks the residue present in H8 undergoing the transition, as described through our simulation data. Based on our experiment and simulation data, we have been able to associate coil-helix transition exchange rates and both forward and reverse kinetics of helix-coil in H8, respectively (C). The asymmetric exchange kinetics between the forward (helix to coil) and reverse (coil to helix) pathways from simulations (in black) whereas, from E-CPMG NMR (in blue) we were able to determine the reverse (coil to helix) path only which shows overlap with the MD simulation (a difference of ∼19%). Apart from the timescales, we also highlight the percentage of helix and coil populations from both NMR and MD simulations.

Equilibrium flux analysis reveals pronounced kinetic asymmetry between these basins (Fig. 3A). Exchange within the coil ensemble is rapid and highly interconnected, with transitions between M0 and M4 occurring on microsecond timescales (Supplementary Table 4). In contrast, transitions toward the fully helical state proceed predominantly through the intermediate ensemble and occur substantially more slowly. Direct transitions from the coil states into M3 are rarely observed, indicating that complete propagation into a 10-residue helix represents a kinetically disfavored event. Consistent with this interpretation, MSM-derived mean first passage times^49^ show rapid helix-to-coil relaxation (MFPT _HtoC_ ≈ 2-4 μs) but markedly slower coil-to-helix transitions (MFPT_CtoH_ ≈ 20–45 μs) across the validated MSM models (Fig. 3C). Model validation demonstrated convergence of the slowest implied timescales, Chapman-Kolmogorov^47^ behavior and mean first passage times across lag times and cluster numbers, supporting the robustness of the six-state kinetic description (Supplementary Fig. 7).

The presence of fast conformational exchange of H8 was independently examined using extreme-CPMG (E-CPMG)^45^ relaxation dispersion experiments at 272K. It implicates residues in H8 showing very fast kinetics of exchange (Fig. 3B; Supplementary Fig. 8,10). Residues within the central hydrophobic segment of H8 (151-153) exhibit clear exchange broadening consistent with microsecond-timescale conformational exchange, whereas the acidic patch (147-149) and K155 show little detectable exchange (Supplementary Table 5). The similarity in the exchange behaviour across residues 151-153 corroborates with frequent nucleation in this region, while the heterogeneous residue participation across H8 further supports weak cooperativity during helix propagation.

Exchange was not detected at higher or lower temperatures (262 K, 277 K and 282 K; Supplementary Fig. 9), indicating that the conformational exchange occurs within a narrow experimentally accessible timescale window. E-CPMG measurements estimate a global exchange timescale of ∼25 μs for H8 residues (Fig. 3B), consistent with the slower coil-to-helix transitions predicted by MSM. Although the MSM resolves multiple intermediate macrostates whereas E-CPMG reports an effective exchange process between NMR- distinguishable conformations, both approaches independently support the presence of low-population transient helical states undergoing microsecond-timescale exchange. Consistent with this interpretation, MSM analysis predicts that the fully helical macrostate (M3) constitutes only ∼0.1% of the equilibrium population, whereas the broader partially helical intermediate ensemble (M1, M2 and M5) collectively accounts for ∼5% of the conformational landscape. The remaining ∼95% population resides within the coil basin (M0 and M4) (Supplementary Fig. 7). Beyond H8, several flexible loop regions (in Loop1, Loop2 and V123, A124, R136) in Skp1 also exhibit enhanced exchange behaviour in the E-CPMG experiments, reflecting the broader conformational dynamics of the protein (Supplementary Fig. 8).

### Residue-specific interactions stabilise transient helices in H8

The weak cooperativity underlying helix formation in H8 suggests that local residue-residue interactions are insufficient to support propagation to a fully extended helix. To identify interactions associated with helix nucleation and stabilisation of transient conformations, we compared residue-level contact maps from single trajectories that represent H8 in helical conformation (Fig. 4A), intermediate (Fig. 4B) and coil (Fig. 4C) conformations, respectively. The coil conformation exhibits broadly distributed contacts between H8 and distal regions of Skp1, including interactions extending toward H5 and H6 (Fig. 4C). In contrast, the helical conformation is characterised by dense intra-H8 interactions, indicating local compaction and increased contact cooperativity within the segment (Fig. 4A). The intermediate conformations retain a subset of these local contacts but lack the extensive interaction network observed in the fully helical state (Fig. 4B), consistent with incomplete propagation of transient helices. Hydrophobic residues within H8 (A151 and V153) contribute prominently to these locally stabilised interactions in the helical ensemble but are substantially less connected in the coil and intermediate states. The progressive increase in intra-H8 contact density from coil to intermediate to helix suggests that local enthalpic interactions are required to stabilise transient helices and support further propagation. However, the reduced contact density within the intermediate ensemble indicates that these interactions remain insufficient to maintain a persistent helical conformation.

**Figure 4:**
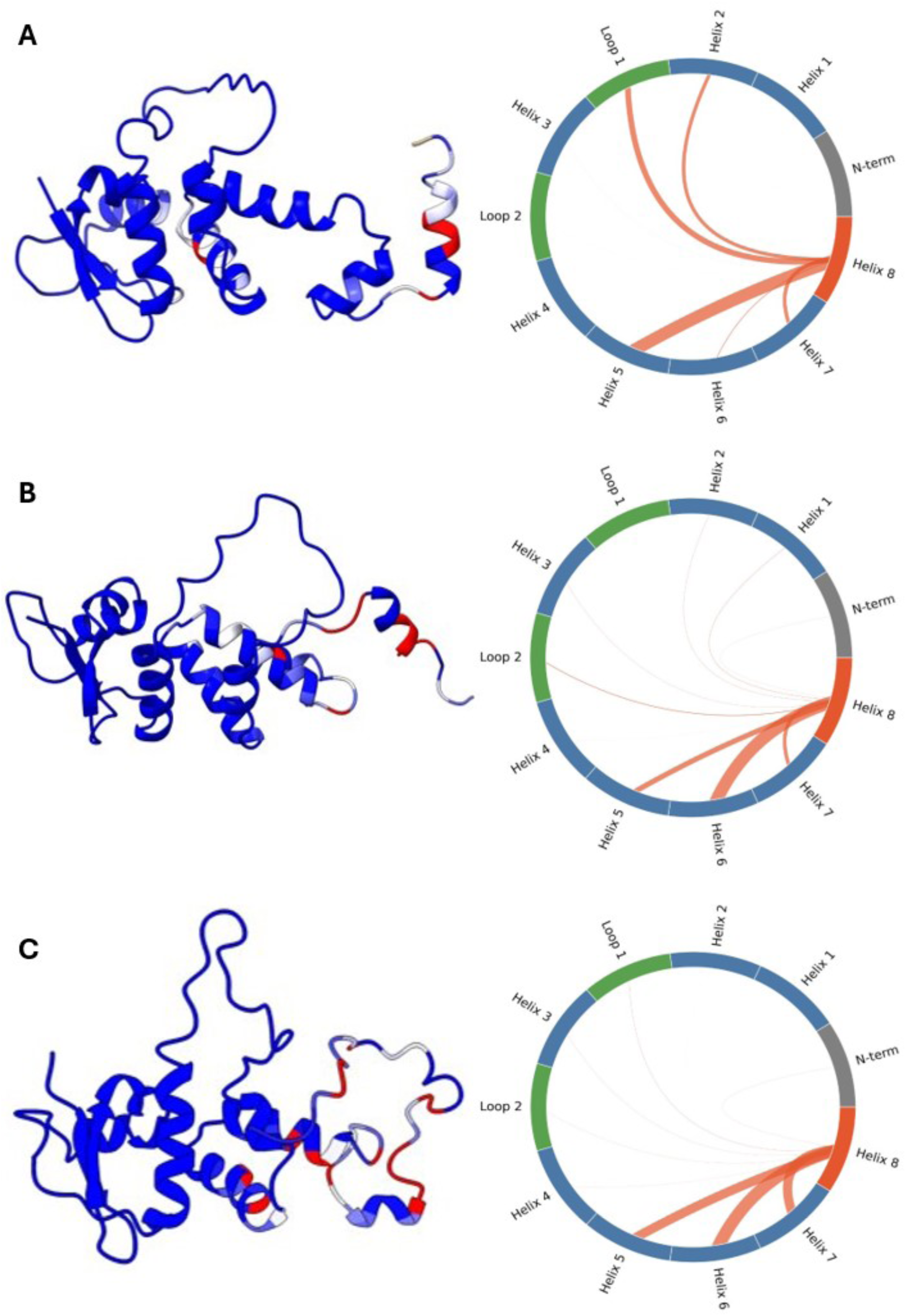
Contact maps generated from specific trajectories of our MD dataset to highlight the low dependence of H8 residues in forming contacts with other segments or regions of the protein in helical state than when compared to the coil state highlights a key dependence on intra-local interactions within H8 residues for the “order” state. Chord wheel diagrams highlight the key interactions between H8 residues (147-156) in making frequent contacts with other regions of the protein (thicker the width of the curve associating H8 with other regions, the more frequent the contact). Figures A-C are described in a descending manner of change in ordering or helicity in H8. Regions highlighted in red across Skp1 points to the regions where highly frequent contacts are made.

The presence of both negatively and positively charged residues within H8 further suggests a role for electrostatic interactions in transient helix stabilisation. Interactions between the acidic patch and R154 are enriched in the helical and intermediate ensembles (Supplementary Fig. 11). During helix unfolding, these interactions weaken as the residue separation increases beyond ∼5 Å, whereas transient helix formation is associated with closer approach (∼2 Å), consistent with transient salt-bridge formation during local helix stabilisation.

### Protonation of the acidic patch promotes helix formation in H8

Previous analyses indicated that helix formation within H8 is driven primarily by local intra-segment interactions and remains restricted to short partially propagated helices. To determine whether these transitions are intrinsically encoded within H8, independent of the remainder of Skp1, we synthesised an isolated peptide spanning residues 144-163 and examined its conformational behaviour using circular dichroism (CD) spectroscopy. Helix formation within the isolated H8 peptide exhibits strong pH dependence (Supplementary Fig. 12A), with maximal helicity observed at pH 3.4 (fractional helicity of 0.4) and progressively reduced helicity toward neutral pH (Fig. 5A). Thermal denaturation of H8 (Fig. 5B) similarly reveals increased helix stability at low pH, with the peptide at pH 3.4 displaying an increase in melting temperature (T_m_) of ∼12°C relative to pH 7 (Supplementary Fig. 12B and 13). These observations demonstrate that H8 alone is sufficient to undergo helix-coil transitions and that helix stability is strongly modulated by protonation state. The glutamate-rich acidic patch within H8 is highly negatively charged at neutral pH. Protonation of these glutamate residues at lower pH reduces electrostatic repulsion within the segment and promotes local packing interactions required for helix extension. Consistent with this interpretation, the helical ensemble exhibits increased intra-H8 contacts and transient salt-bridge formation involving R154, interactions that are substantially weakened in the coil ensemble. Neutralization of the acidic patch therefore favours propagation beyond short locally stabilized helices.

**Figure 5:**
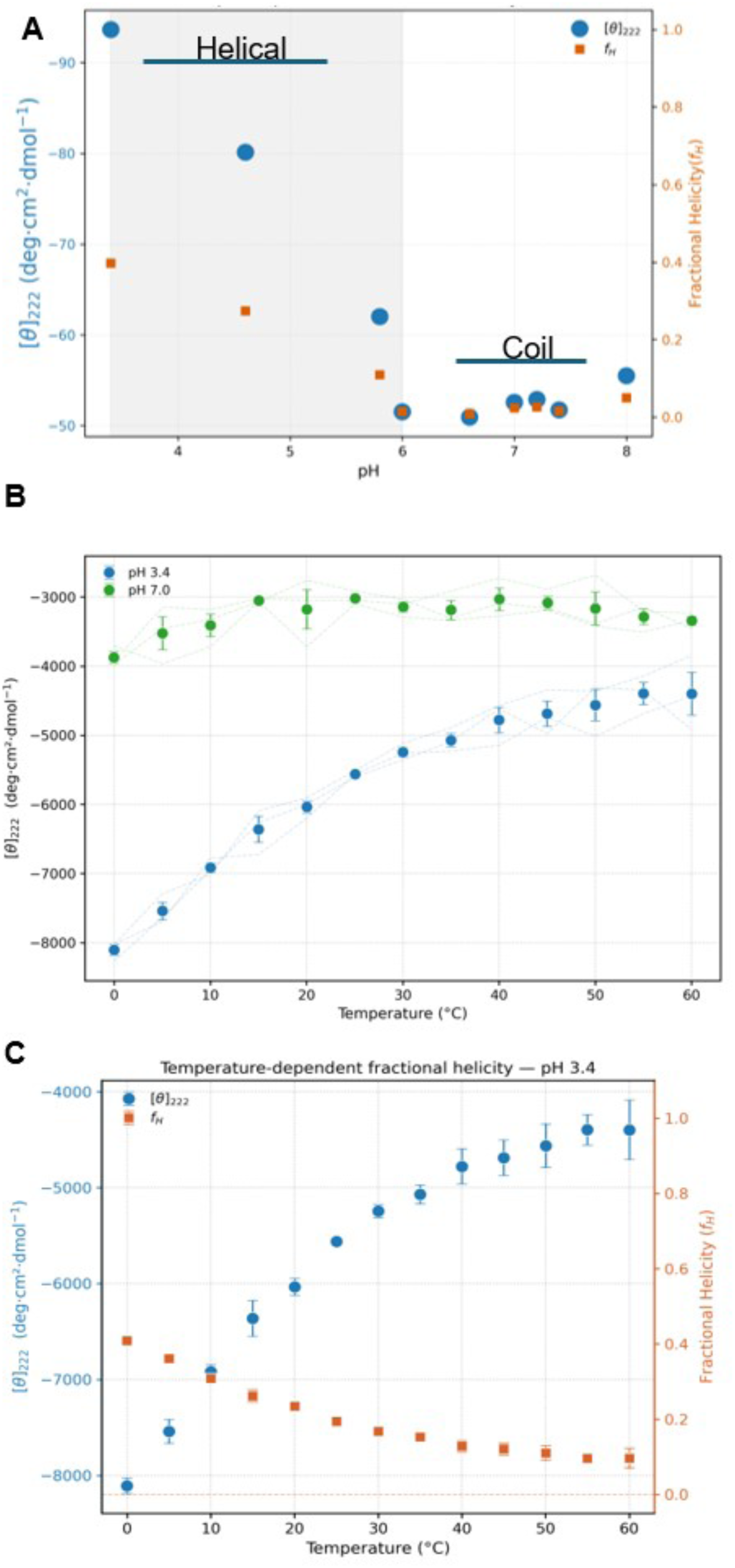
CD experiments carried out on H8 peptide to determine transient helicity as a function of pH. Mean residual ellipticity at θ_222_ measured from CD (blue circles) was compared against a range of pH (8-3.4) (A) to show that lowering of pH leads to an increase in fractional helicity (yellow squares) with the maximum helicity of 0.4 at pH 3.4. Shaded region in (A) highlights increase in fractional helicity at pH <6. Based on this, the two extremes, one at maximum fractional helicity, and one at minimum fractional helicity were compared through a thermal melt. The fractional helicity observed at pH=3.4 was shown to have a significant thermal denaturation owing to the presence of residual helicity as compared to the profile at pH=7(B). In both cases, the thermal profile ranges from a T=0-60°C. Standard errors of mean were calculated for MRE observed at θ_222_. The thermal profile at pH=3.4 was used for determining change in fractional helicity(C).

**Figure 6:**
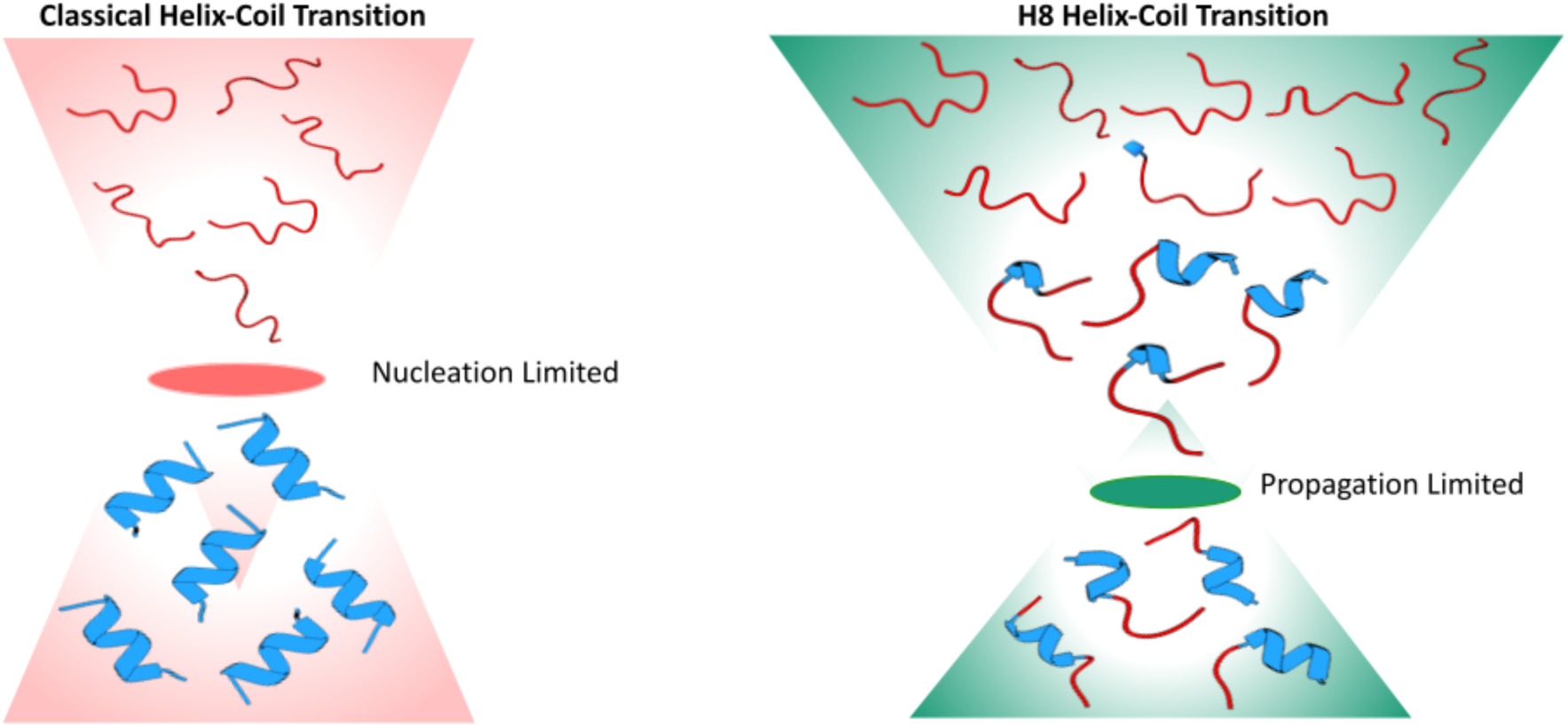
Distinct bottlenecks that defines underlying classical and H8 helix-coil transitions are represented through a classical helix-coil transition, where helix formation is primarily nucleation limited and the major energetic barrier lies in the initial formation of a stable helical nucleus from the disordered ensemble (Left). Once nucleated, helix propagation proceeds cooperatively to generate fully helical conformations. In contrast, the H8 helix-coil transition is predominantly propagation limited (Right). Transient local helical segments readily nucleate within the disordered ensemble, but sequence- encoded constraints prevent efficient propagation into a continuous stable helix, resulting in heterogeneous partially folded intermediates and a frustrated conformational landscape.

Despite this stabilization, the isolated peptide does not adopt a persistent fully helical conformation, indicating that protonation shifts the equilibrium toward increased helicity without eliminating the underlying dynamic exchange behavior. The calculated free energy change for the helix–coil transition (∼-2 kcal/mol at physiological temperature) is consistent with previous measurements^50^ of transient helical systems. These observations provide a physical basis for the weak cooperativity observed in H8. Frequent local nucleation events occur across the segment, but extended helix propagation remains limited under physiological conditions. Rather than stabilising a persistent, folded state, the acidic patch appears to maintain H8 within a dynamically exchangeable ensemble that can be transiently stabilised under favourable electrostatic conditions. Such regulation may prevent kinetic trapping in a single helical conformation and preserve the conformational plasticity required for interaction with the diverse FBPs.

## Discussion

Skp1 H8 provides a tractable system for understanding how transient secondary structure is encoded within an IDR involved in molecular recognition. In the F-box bound state, H8 is in a helical conformation^37^, whereas in apo-Skp1 the same region is predominantly disordered^36^. Using an interdisciplinary strategy that integrates atomistic simulations, kinetic modelling, relaxation dispersion NMR and peptide biophysics, we show that H8 does not undergo a simple two-state helix-coil transition. Instead, the segment populates a heterogeneous coil-dominated ensemble containing short-lived partially helical intermediates that repeatedly form but rarely propagate into a persistent full-length helix.

A central finding of this work is that helix formation within H8 is propagation-limited rather than being nucleation-limited. Classical helix-coil models generally describe nucleation as the dominant energetic barrier followed by comparatively favourable propagation^30,31^. H8 exhibits the opposite behaviour. These models have largely been developed from peptide systems in which helicity is either directly induced or already substantially populated, making helix unfolding easier to describe than helix formation from a coil-dominated IDR ensemble. H8 presents a different context: a predominantly disordered segment that repeatedly nucleates short helical elements, either as pre-existing conformers or as states available for partner stabilisation but rarely propagates into a persistent helix in the apo-state. Local nucleation appears readily accessible from multiple positions within H8, consistent with the distributed intrinsic helix-forming propensity of several residues. However, the sequence grammar appears tuned to prevent persistent helix propagation and preserve conformational plasticity for FBP recognition. Consequently, the conformational landscape is dominated by partially propagated helices rather than a clear bimodal equilibrium between coil and helix states.

The kinetic asymmetry observed for H8 is consistent with a broader pattern in which low-population helical states are accessible but thermodynamically disfavoured^20^. Ultrafast peptide studies typically place local helix-coil motions in the nanosecond-to-sub-microsecond regime^51,52^, whereas relaxation-dispersion NMR resolves slower exchange involving sparsely populated excited states^45^. Comparable behaviour has been reported for systems such as the PBX homeodomain C-terminal extension, where a minor helical state exchanges with a predominantly disordered ensemble on the low-microsecond timescale^53^, and for the KIX domain, where helix-forming transitions are substantially slower than helix dissolution^54^.

In Skp1, this exchange was detectable only by E-CPMG at 272 K; at higher or lower temperatures, no resolvable relaxation-dispersion signal was observed, suggesting that cooling shifts H8 dynamics into the experimentally accessible exchange window^55^. Although the simulations and E-CPMG experiments were performed at different temperatures and report different observables, they converge on the same mechanistic picture: H8 samples low-population transient helical states undergoing microsecond-timescale exchange, with productive helix formation being slower than helix loss. H8 therefore follows the general kinetic asymmetry observed in other transient helical systems but differs in its underlying mechanism: the transient helical ensemble is not limited by lack of nucleation, but by inefficient propagation from multiple locally nucleated segments.

The glutamate-rich acidic patch plays a central role in regulating this behaviour in H8. Protonation of the isolated H8 peptide substantially increases helicity and thermal stability, demonstrating that H8 intrinsically encodes helix-forming capacity and that this capacity is highly sensitive to charge state. Under physiological conditions, clustered glutamates likely limit persistent helix propagation through electrostatic repulsion between neighbouring backbone dipoles and side chains, while still permitting transient local nucleation. This acidic patch should not be viewed as a simple destabilising feature, with protonated glutamate residues being identified as having highest helix propensity after alanine^56^. We propose that it functions as a charge-sensitive conformational rheostat that maintains H8 within a dynamically exchangeable ensemble. This is consistent with emerging evidence that charge composition and spatial patterning of residues within IDRs directly regulate structural propensity and conformational dynamics^57^. F-box binding may overcome this propagation ceiling by stabilising contacts that are only transiently sampled in the apo-ensemble. Polyampholyte theory and charge-patterning analyses have established that the linear organisation of charged residues controls compaction and structural bias within disordered regions^58^. Such regulation may prevent kinetic trapping in a single helical conformation and preserve the conformational plasticity required for interaction with structurally diverse FBPs. The broader occurrence of similarly organised acidic-hydrophobic-basic motifs across disordered regions of the proteome (Supplementary Fig. 15) further suggests that propagation-limited transient helicity may represent a more general mechanism for regulating conformational exchange and molecular recognition.

This places H8 within an α-MoRF(Molecular Recognition Feature)^6^ like recognition regime, but with a distinct feature: unlike MoRFs dominated either by partner-induced folding or substantial pre-organisation, H8 remains strongly coil-dominated, with repeated helix nucleation and restricted propagation. Full propagation into the bound helical state therefore likely requires partner-specific stabilisation(Supplementary Figure 16). This interpretation is consistent with the structural diversity observed across Skp1 F-box complexes, where the extent of helicity within H8 varies between bound conformations(Supplementary Figure 17).

This study also has limitations that define clear next steps. Although the two simulation ensembles capture repeated helix-coil interconversion, the number of complete coil-to-helix and helix-to-coil transitions remains limited relative to the complexity of the H8 landscape, and the MSM-derived rates should therefore be interpreted as effective kinetic estimates rather than exhaustive sampling of all pathways. Experimentally, multi-nuclear chemical shift measurements and additional relaxation-dispersion datasets would further refine residue-level assignments of the transient helical ensemble.

To conclude, H8 illustrates how an IDR can encode conditional structure without committing to a stable fold. Its charged-hydrophobic-charged architecture permits local helix nucleation while limiting long-range propagation, creating a metastable recognition element that preserves conformational flexibility. This mechanism suggests a design principle for engineering MoRF- like segments with tunable structural propensity, where charge density and spacing around helix-prone motifs modulate the balance between conformational plasticity and partner- stabilised structure.

## Materials and Methods

### MD Simulation

Initial coordinates for molecular dynamics (MD) simulations were derived from two Skp1 structures representing distinct H8 conformations: the F-box-bound crystal structure (PDB: 1FQV), in which H8 is helical, and the apo solution NMR structure (PDB: 5XYL), in which H8 is disordered, as described previously^37^. These systems are referred to as Skp1_helix_ and Skp1_coil_ respectively. All simulations were performed using GROMACS (v.2022)^59^. The protein was described using the Charmm36m force field^60^, and water molecules were modelled using the TIP3P water model^61^.

Each protein structure was placed in a dodecahedral simulation box with a minimum distance of 1.25 nm between the protein and the box boundary. The system was solvated with TIP3P water and neutralised with 150 mM NaCl. Energy minimisation was performed using the steepest descent algorithm until the maximum force was below 1000 kJ mol⁻¹ nm⁻¹. Temperature equilibration was carried out at 300 K for 100 ps using the velocity-rescale thermostat ^62^ with a coupling constant of 0.2 ps, followed by pressure equilibration for 500 ps at 1 atm using the Berendsen barostat ^63^ with a coupling constant of 1 ps.

Production simulations were performed with temperature maintained at 300 K using the velocity-rescale thermostat with a coupling constant of 0.1 ps and pressure maintained at 1 atm using the Parrinello–Rahman barostat^64^ with a coupling constant of 2.0 ps^65^. Bonds involving hydrogen atoms were constrained using the LINCS algorithm^66^. Coulombic and van der Waals interactions were treated with a 1.2 nm cut-off. Trajectory coordinates were saved every 20 ps. Simulation lengths for all replicas are provided in Supplementary Tables 1 and 2.

### Helix Formation and Analysis

Skp1_helix_ and Skp1_coil_ were analysed to quantify H8 helicity and characterise transient helix formation using DSSPv3.0^67,68^. Secondary-structure assignments were obtained at 100 ps intervals. Eight DSSP states were considered: coil (∼), β-sheet (E), β-bridge (B), bend (S), turn (T), α-helix (H), π-helix (I) and 3_10_-helix (G). For subsequent analysis, these states were reduced to a two-state representation. Residues assigned as H, I or G were classified as helical, whereas residues assigned as coil, β-sheet, β-bridge, bend or turn were classified as non-helical. Transient helices were defined as conformations containing at least five contiguous helical residues within H8 (residues 147-156), thereby excluding discontinuous or sparsely populated helical configurations. To ensure persistence, only helical segments maintained for more than 20 consecutive frames (∼2 ns) were considered transient helix events. For Skp1_helix_ trajectories the initially folded H8 segment was required to unfold first, defined as fewer than five helical residues within H8, before subsequent transient helix events were included in the analysis. Representative conformations corresponding to helices of varying lengths shown in Fig. 2A-B were extracted from MD trajectories and rendered using UCSF ChimeraX^69^.

### Markov state models

C_⍶_ coordinates corresponding to the H8 segment (residues 147-156) from Skp1_helix_ and Skp1_coil_ trajectories were used for feature construction. A C_⍶_ based helix score and writhe^48^ descriptors were employed to quantify local helical geometry and conformational topology. Features from both ensembles were pooled and subjected to time-lagged canonical correlation analysis (tCCA)^70^ to identify slow collective coordinates associated with helix-coil exchange. The leading tCCA component together with the helix score was subsequently used for k-means clustering^71^ to define discrete microstates.

Transition probability matrices were estimated across varying lag times and symmetrized to enforce detailed balance. Markovian behavior was evaluated through implied timescale convergence and Chapman-Kolmogorov validation tests^46,47^. Mean first-passage times (MFPTs) and interconversion rates between kinetically dominant states were calculated from the transition matrix to quantify asymmetric helix-coil exchange dynamics within H8. The effective transition rate from state (i) to state (j) was defined as:

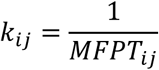

Net reactive flux between states was further computed using the stationary distribution (πᵢ) and transition probabilities (T_ij_) to estimate equilibrium probability currents and identify dominant kinetic pathways associated with helix-to-coil and coil-to-helix exchange:

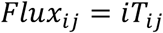

### Contact Analysis

Residue–residue contacts involving H8 residues (147-156) were computed based on minimum interatomic distances. Contacts were evaluated every fifth simulation frame and were defined by any pair of atoms within 4.5 Å. To exclude local sequence effects, residue pairs satisfying (|i-j| <3) were omitted from the analysis. Contact frequency for each residue pair was calculated as the fraction of analysed frames containing at least one interatomic contact. In parallel, a scaled contact metric was determined by normalizing the number of contacting atom pairs against the maximum possible atom-atom contacts for the corresponding residue pair. Only contacts exhibiting persistence greater than 25% were retained for visualization. Symmetric contact matrices were generated and represented as heatmaps, with interaction intensities proportional to contact frequencies.

### Skp1 expression and purification

Skp1 gene construct has been inserted into the pGEX-6p-1 vector between BamH1 and Xho1 sites. The Skp1-pGEX-6p-1 plasmid construct was transformed in E. coli BL21 DE3 cells and was grown in M9 minimal media supplemented with ^13^C-glucose and/or ^15^N–NH_4_Cl (Cambridge Isotope Laboratories, Andover, MA and Eurisotop, France) for up to 0.6–0.8 O.D. (A600) at 37 °C temperature. The Skp1 fusion protein expression was induced overnight by adding 200 µM isopropyl β-D-1-thiogalactopyranoside (IPTG) at 25 °C, and the induced culture was harvested. The cells were suspended in sodium phosphate buffer (20 mM, pH 6.0), 1 mM EDTA, 100 mM NaCl, 1 mM β-ME, 1 mg/ml lysozyme, 0.1% Triton X-100 followed by sonication for cell lysis. The cell lysate was bound to GSH-Sepharose beads, and the beads were washed with a sodium phosphate buffer with an increasing NaCl gradient. The GST-tag was removed by incubating with PreScission protease at 4 °C overnight. The protocol is similar to used in our previous work^38^.

### E-CPMG Relaxation Dispersion NMR Experiments

All NMR experiments were performed using ^15^N-labelled Skp1 (similarly prepared as discussed above). Data were acquired on Bruker 800, 900, and 950 MHz spectrometers equipped with cryogenic probeheads optimized for high-power E-CPMG measurements^45^. In the E-CPMG experiments, refocusing pulses were applied using a high-power ^15^N B(1) field of 6 kHz across all refocusing frequencies. Backbone amide ^15^N E-CPMG relaxation-dispersion experiments were recorded at 277 K, 272 K, and 262 K. To maintain the sample in the liquid state under near-freezing conditions, samples were loaded into capillary tubes, which minimized freezing during low-temperature measurements. Specialized heavy NMR spinners were additionally used to stabilize the lightweight capillary assemblies during spectrometer insertion and data acquisition. All experiments were recorded with a recycle delay of 1.5 s. The CPMG relaxation period was optimized by testing delays between 10-40 ms and was set to 20 ms for all subsequent experiments. NMR data were processed using NMRPipe^72^ and analyzed in CARA^73^. Peak lists, peak tables, and intensity data exported from CARA were subsequently analyzed in Mathematica^74^ to generate (R_2eff_) dispersion profiles as a function of CPMG frequency^45^. Local and global fitting of the relaxation-dispersion data were performed using ShereKhan^75^.

### Peptide synthesis

A 20 amino acid peptide (DFTEEEEAQVRKENQWCEEK) was purchased from GenScript USA, Inc. The N-terminal was acetylated for protection, and its purity was more than 95% (as determined by manufacturer).

### Circular Dichroism

Far-UV Circular Dichroism analyses were performed using stock solutions of peptide at a concentration of 1mg/ml in ultrapure water. For, pH-dependent measurements, peptide solutions were diluted to 40 uM either in either 20 mM sodium acetate buffer with 100 mM sodium chloride for pH ranges of 3.4-5 or in phosphate buffer, 20 mM with 100 mM sodium chloride for pH ranges of 5.8-8. Temperature dependent experiments were carried out at pH 3.4 and 7.4, across a temperature range of 0 °C to 50 °C, in 1 °C increments. Spectra were acquired using a Jasco J-1500 spectropolarimeter (Jasco Corp., Tokyo, Japan) equipped with a Peltier temperature control system and a 1mm quartz cuvette. Measurements were collected at 222 nm, at a scanning speed of 100 nm/min, with a data integration time (DIT) of 4 seconds and 3 accumulations per sample. Fractional helicity at each pH and temperature was calculated^76^. Thermal melts of the samples were fitted using Van’t Hoff equation to estimate melting temperatures (T_m_) and associated change in free energy (ΔG)^77^.

