## Supplementary Information for "A Charge-Encoded Rheostat Permits Helix Nucleation but Limits Propagation in Skp1"

***Corresponding authors**

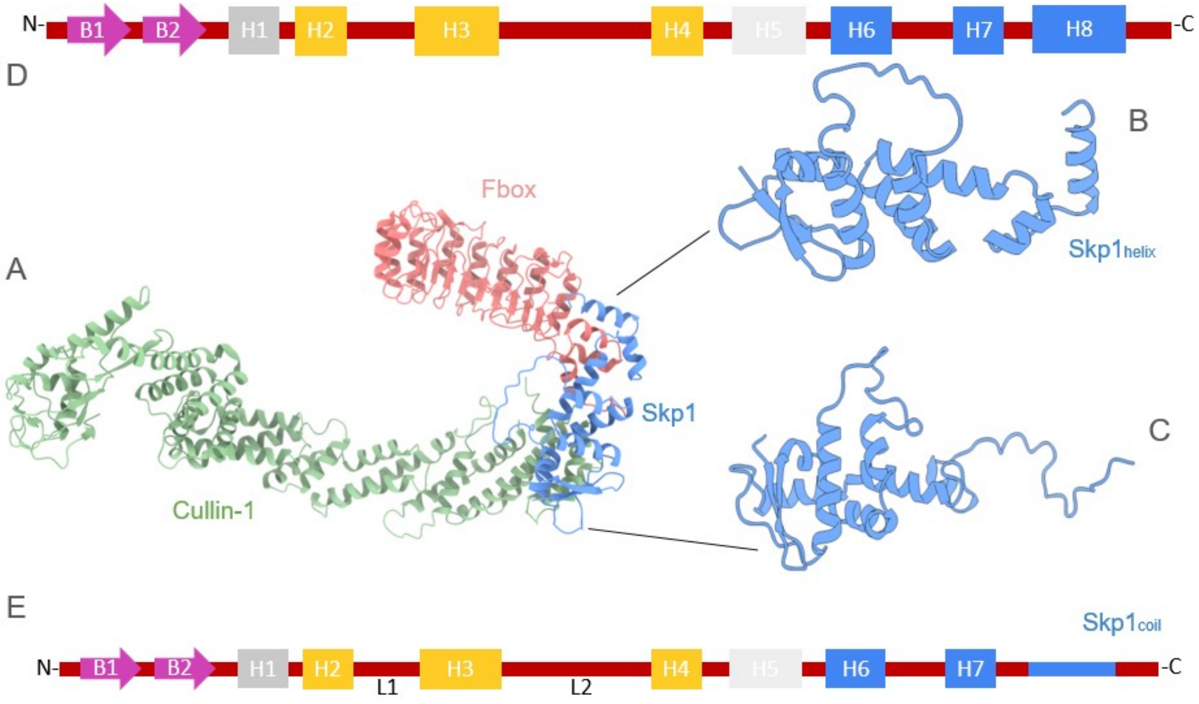


**Supplementary Figure 1: Structural organisation of the SCF E3 ubiquitin ligase complex**. (A)Skp1 (blue) occupies a central adaptor position and forms two distinct interaction interfaces: one with Cullin1 (green) and another with the F-box protein Skp2 (salmon). The C-terminal helices H6–H8 of Skp1 mediate recognition of the F-box domain, whereas a separate interface interacts with Cullin1. The $Skp1_{helix}$ structure (left) was derived from the F-box-bound crystal structure (PDB: 1FQV) (B), whereas the $Skp1_{coil}$ structure (right) was derived from the apo NMR structure (PDB: 5XYL) (C). The principal differences are localised to the C-terminal region encompassing helices H6–H8, with H8 adopting a fully helical conformation in Skp1helix and a disordered conformation in Skp1coil (D,E).

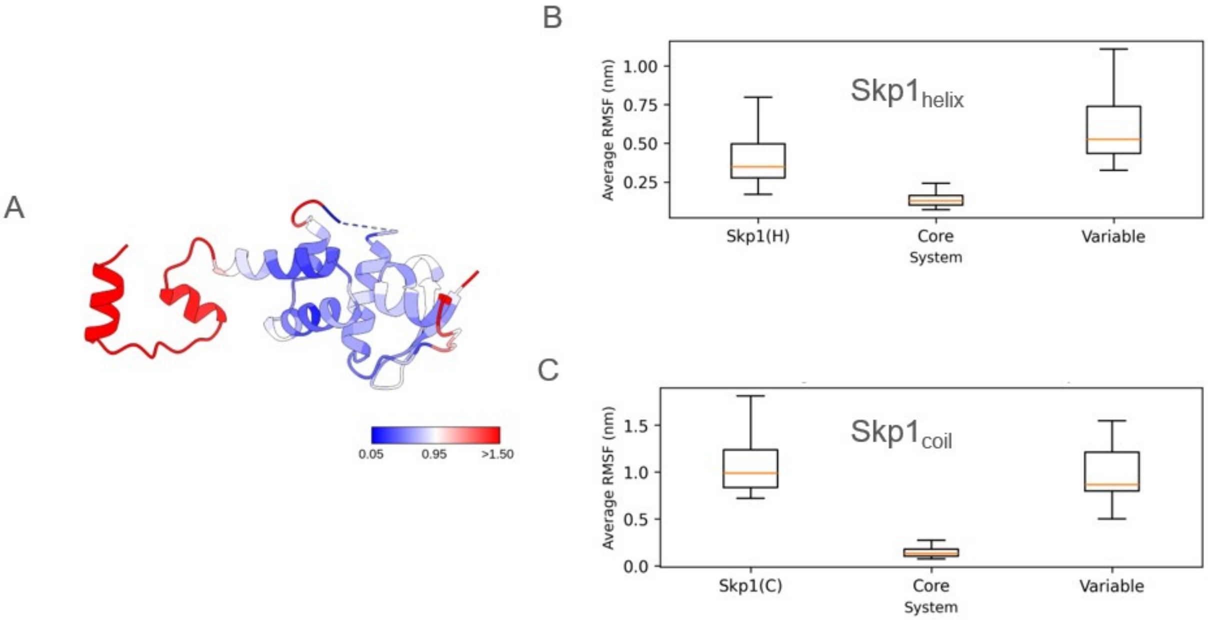


**Supplementary Figure 2: Structural variability and residue-level flexibility across Skp1**. (A) Structural variability estimated from RMSD across experimentally determined Skp1 structures listed in Supplementary Table 7. Regions with higher variability (>1.5 Å) are highlighted in red. The structurally stable core comprises helices H1–H5, whereas the variable region includes Loop1, Loop2 and the C-terminal H6–H8 segment. (B–C) Average residue-level RMSF calculated from the Skp1helix and Skp1coil simulation ensembles. In both ensembles, fluctuations are largely restricted to the variable regions, while the Skp1 core remains comparatively rigid, with average RMSF values below 0.5 nm.

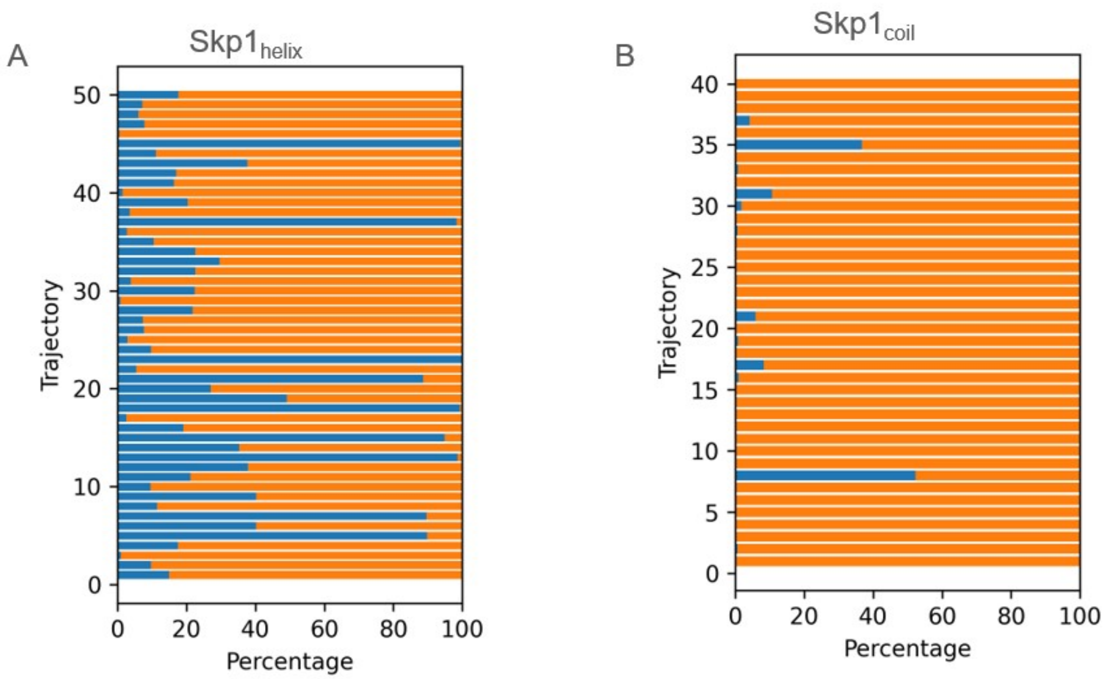


**Supplementary Figure 3: H8 populates a predominantly coil-dominated ensemble with variable transient helicity.** Distribution of helical and coil populations within H8 (residues 147–156) across individual trajectories of the $Skp1_{helix}$ (A) and $Skp1_{coil}$ (B) ensembles. The percentage of simulation time spent in helical (blue) and coil (yellow) conformations was calculated from DSSP secondary-structure assignments. Both ensembles are dominated by coil conformations, although transient helical populations are sampled in a subset of trajectories.

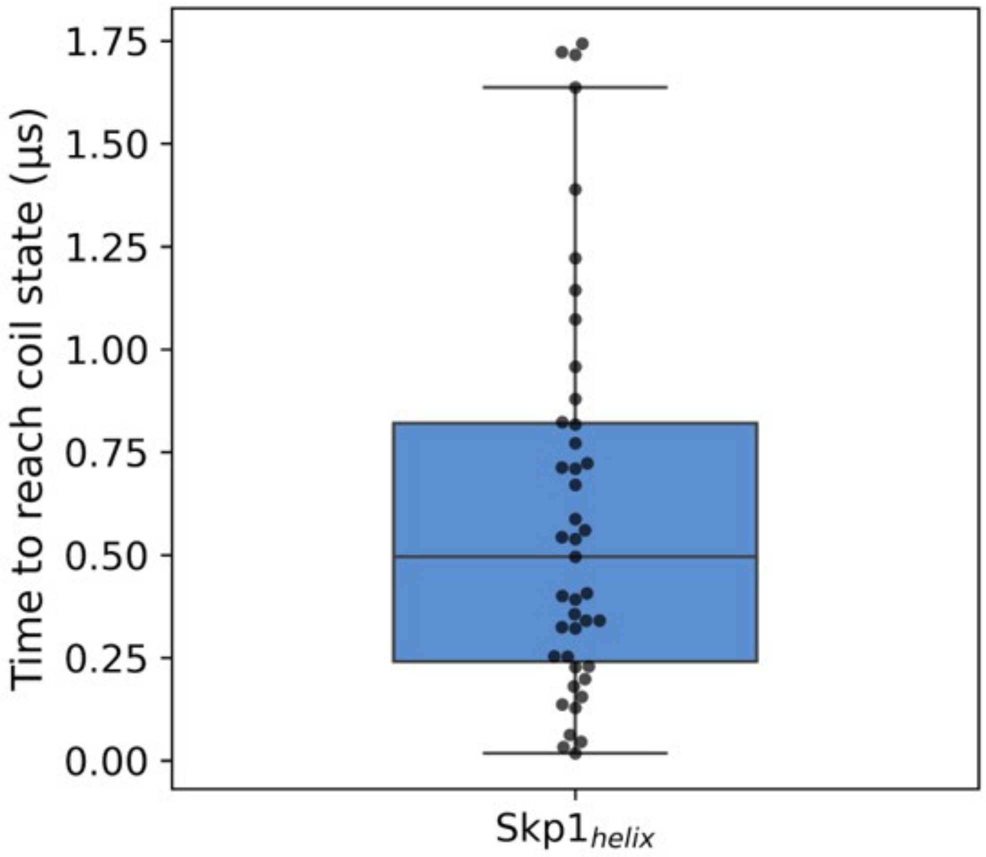


**Supplementary Figure 4: H8 exhibits heterogeneous helix unfolding kinetics across the Skp1helix ensemble.** Distribution of helix persistence times in the $Skp1_{helix}$ ensemble. As H8 is initially helical in all $Skp1_{helix}$ trajectories, this distribution reflects the time required for loss of the starting helical conformation. Helix persistence varies substantially across trajectories, ranging from less than 0.25 μs to approximately 1.75 μs, indicating heterogeneous unfolding kinetics within the ensemble.

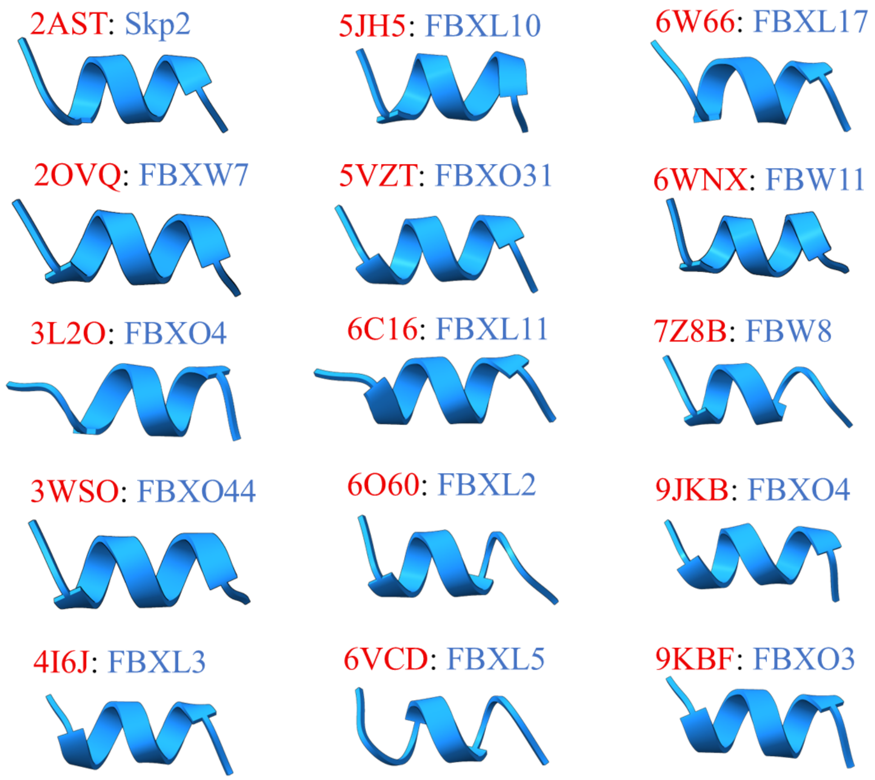


**Supplementary Figure 5: Variability in Helix 8 of Skp1 across different Fbox bound complexes from PDB.** Residues of H8 from 147-156 were extracted from Skp1-bound complexes.

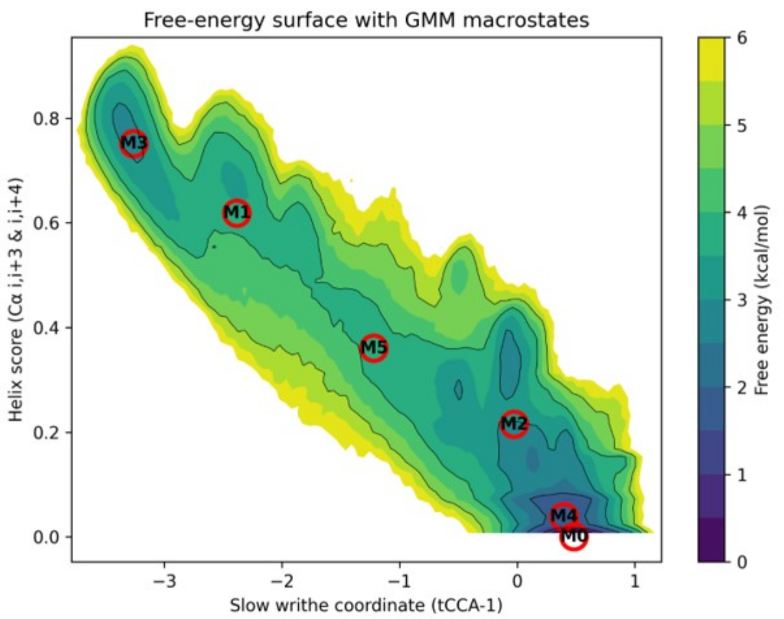


**Supplementary Figure 6: Markov State Model (MSM) resolves the H8 conformational landscape into coil, intermediate and helical macrostates.** Free-energy surface of H8 obtained from MSM using writhe [ref] features extracted from H8 as the reaction coordinate. Six major macrostates are identified, spanning highly populated coil conformations with little or no helicity (M0 and M4), a rare fully propagated 10-residue helical state (M3), and intermediate partially helical states (M1, M2 and M5) that connect the two basins. The distribution of macrostates highlights the dominance of the coil ensemble and the role of intermediate states in mediating helix–coil exchange.

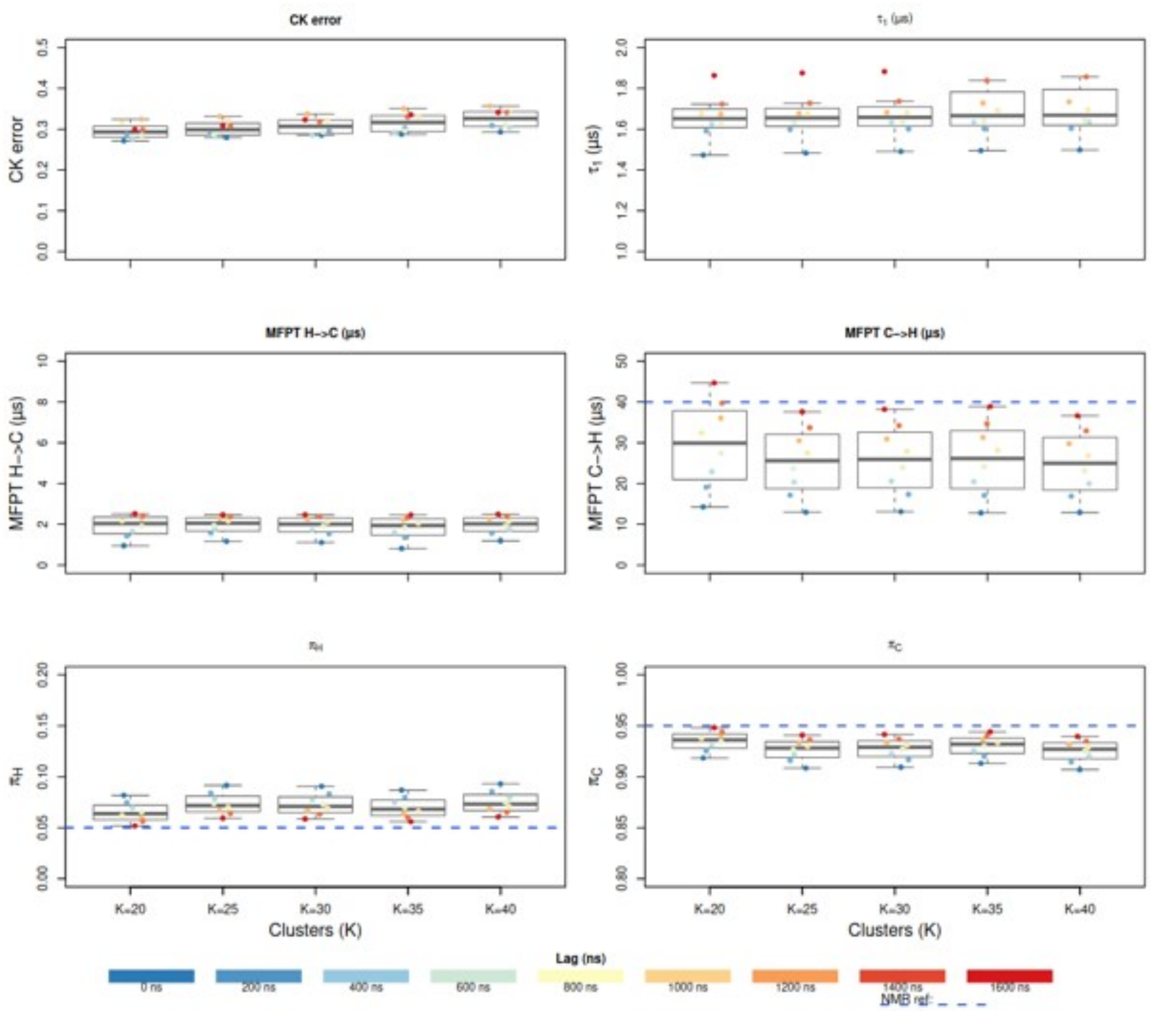


**Supplementary Figure 7: MSM validation of asymmetric helix–coil kinetics in H8.** Boxplots summarising key MSM observables across cluster numbers K = 20–40 and lag times from 0 to 1600 ns. Chapman–Kolmogorov errors remain low across tested models, and the slowest implied timescale converges to ~1.5–1.6 μs, supporting the robustness of the six-state kinetic description. Mean first passage times show rapid helix-to-coil transitions (~2 μs) and slower coil-to-helix transitions (~25–35 μs), consistent with propagation-limited helix formation. The dashed blue line indicates the corresponding E-CPMG-derived exchange estimate (~40 μs). Equilibrium populations of the helical and coil macrostates remain stable across models, confirming a predominantly coil-like ensemble with a low-population helical state. Coloured points represent individual MSM estimates at each lag time, and boxes indicate the interquartile range with median line.

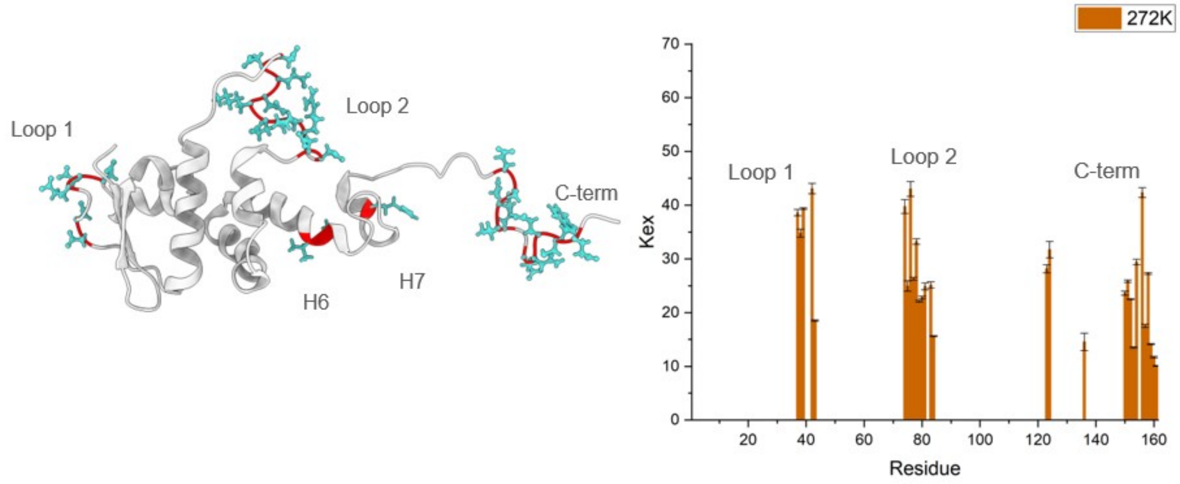


**Supplementary Figure 8: E-CPMG analysis of conformational exchange across Skp1.** Residues exhibiting detectable conformational exchange at 272 K are mapped onto the Skp1 structure (left). Exchanging residues localise predominantly to Loop1, Loop2 and the C-terminal region, with additional isolated residues in H6 and H7. The corresponding exchange rates indicate comparable dynamics across Loop1, Loop2 and the distal C-terminal region. The enrichment of exchange signals in loop and terminal regions is consistent with their enhanced flexibility and their sampling of motions within the E-CPMG-sensitive timescale window.

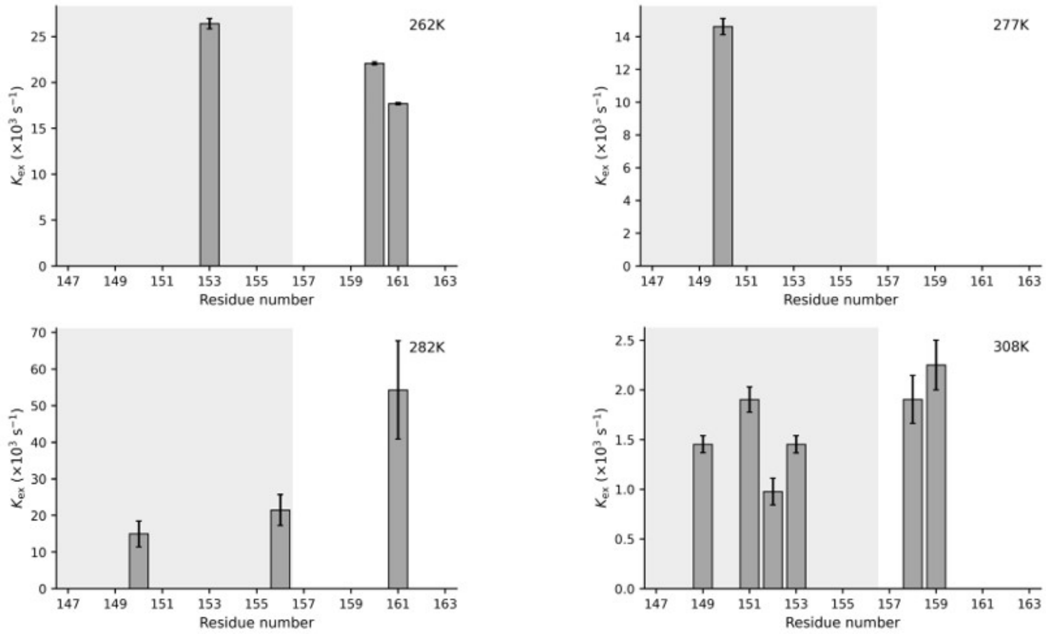


**Supplementary Figure 9: Temperature dependence of C-terminal exchange detected by CPMG/E-CPMG.** $K_{ex}$ values for residues in the Skp1 C-terminal region measured at temperatures other than 272 K. The shaded region marks H8 residues 147–156. Few H8 residues show detectable exchange at these temperatures, in contrast to the exchange observed at 272 K, supporting the use of the 272 K E-CPMG dataset for analysing H8 helix–coil dynamics. At 308 K, conventional CPMG was used.

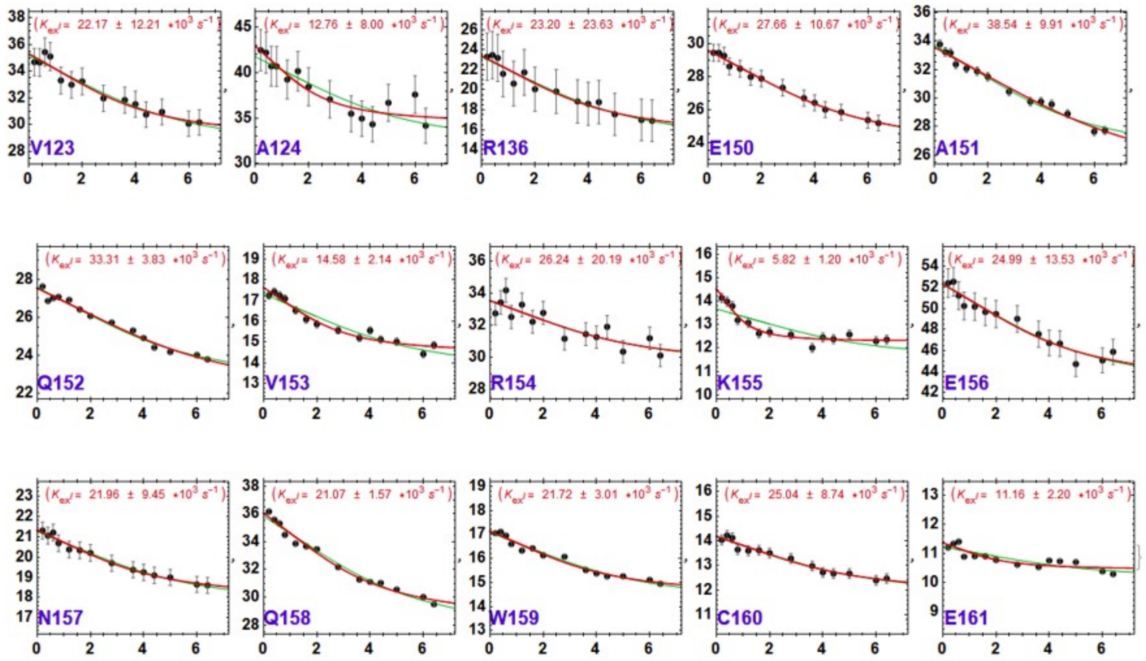


**Supplementary Figure 10: E-CPMG relaxation-dispersion profiles for exchanging residues in the Skp1 C-terminal region.** E-CPMG relaxation-dispersion curves measured at 272 K for residues spanning the C-terminal region of Skp1, excluding V123, A124 and R136. The effective transverse relaxation rate, $R_2$,eff, is plotted as a function of CPMG pulse frequency, νCPMG. Black circles with error bars represent experimental data, and fitted curves are shown in red and green. Extracted exchange rate constants, Kex, are reported in each panel. The magnitude of $R_2$,eff dispersion across the νCPMG range reflects the contribution of conformational exchange, with larger dispersions indicating stronger exchange contributions.

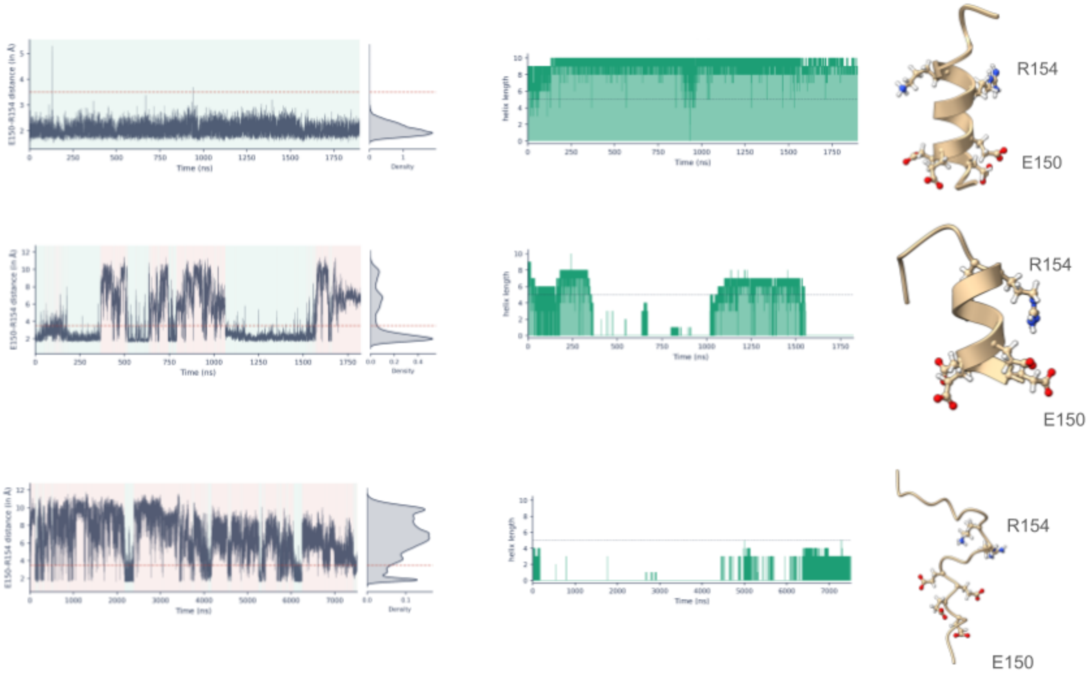


**Supplementary Figure 11: Salt-bridge formation between E150 and R154 correlates with H8 helicity.** Representative structural snapshots (top) and molecular dynamics trajectories from the Skp1$_{helix}$ (top) (extracted from Skp1$_{helix}$ trajectory (23), Skp1$_{intermediate}$ (middle) (extracted from Skp1$_{helix}$ trajectory (9)) and Skp1$_{coil}$ (bottom) (extracted from Skp1$_{coil}$ trajectory). Side chains of E150 (red) and R154 (blue) are highlighted. For each corresponding distance timeseries plot, the change in helix length is also plotted of the trajectory, highlighting that the trajectories are indeed extracted from trajectories that does not undergo any change in helix conformation, undergoes helix-coil transition and includes the intermediate conformation and finally a trajectory that does not reflect any change from coil conformation at all. The helical state is characterised by a persistently short E150–R154 distance (~2 Å), consistent with stable salt-bridge formation, whereas the intermediate and coil states exhibit progressively broader distance distributions indicative of transient or disrupted interactions. The middle panels show the corresponding helix length from DSSP data of the trajectories, with green denoting helical conformations. Sustained helicity is observed in the helical state, frequent helix–coil interconversion in the intermediate state, and predominantly disordered conformations in the coil state.

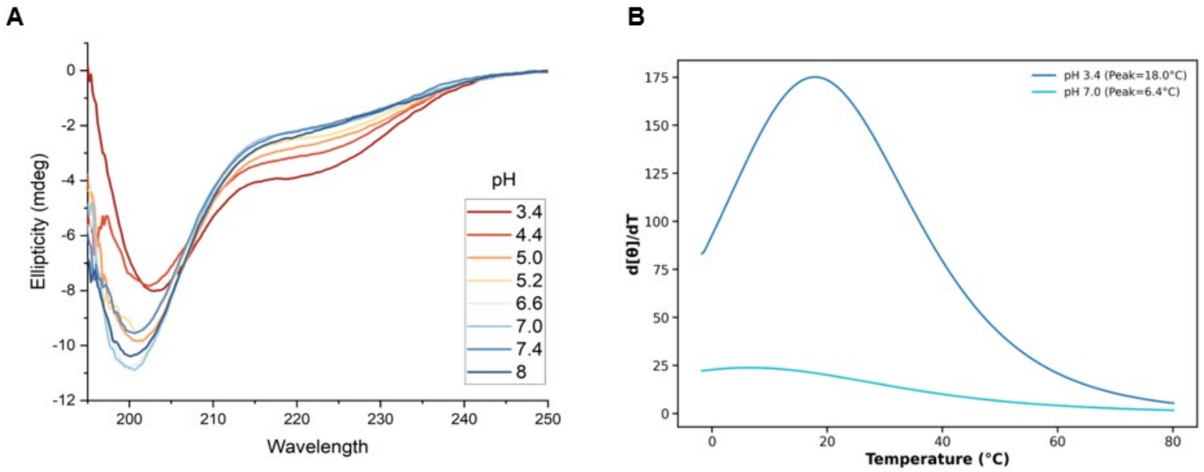


**Supplementary Figure 12: Protonation enhances helicity and thermal stability of the H8 peptide.** (A) Far-UV circular dichroism spectra of the synthetic H8 peptide ($H8_{syn}$) recorded between pH 3.4 and pH 8.0. Spectra exhibit the characteristic double minima at 208 and 222 nm associated with α-helical secondary structure. Helical content increases progressively with decreasing pH, with maximal ellipticity observed at pH 3.4 and reduced helicity at neutral and basic pH values. (B) First derivative of ellipticity with respect to temperature (d[θ]/dT) used to determine the melting temperature ($T_m$) of $H8_{syn}$ at pH 3.4 and pH 7.0. Lowering the pH increases the thermal stability of the peptide, with $T_m$ increasing from 6.4°C at pH 7.0 to 18.0°C at pH 3.4. The larger amplitude of the derivative curve at pH 3.4 is consistent with the greater helical population present under acidic conditions.

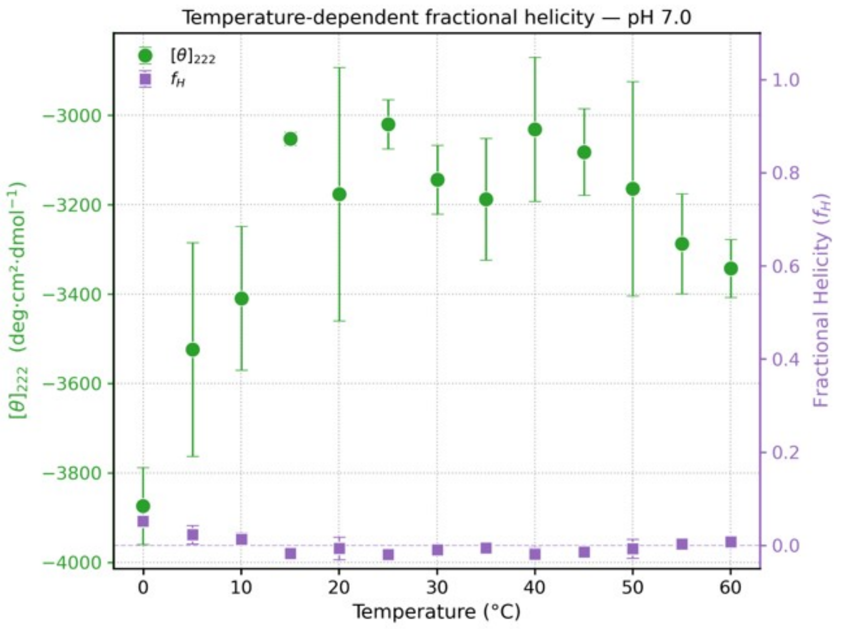


**Supplementary Figure 13: Temperature dependence of residual H8 peptide helicity at neutral pH.** Mean residue ellipticity at 222 nm ($[\theta]_{222}$; green circles, left axis) and calculated fractional helicity (fH; purple squares, right axis) of H8syn plotted as a function of temperature at pH 7.0. $[\theta]_{222}$ values are most negative at low temperature and become progressively less negative with heating, consistent with loss of residual helical structure. Fractional helicity remains very low across the full temperature range, confirming that H8syn is predominantly disordered at neutral pH. Error bars represent standard deviations from replicate measurements.

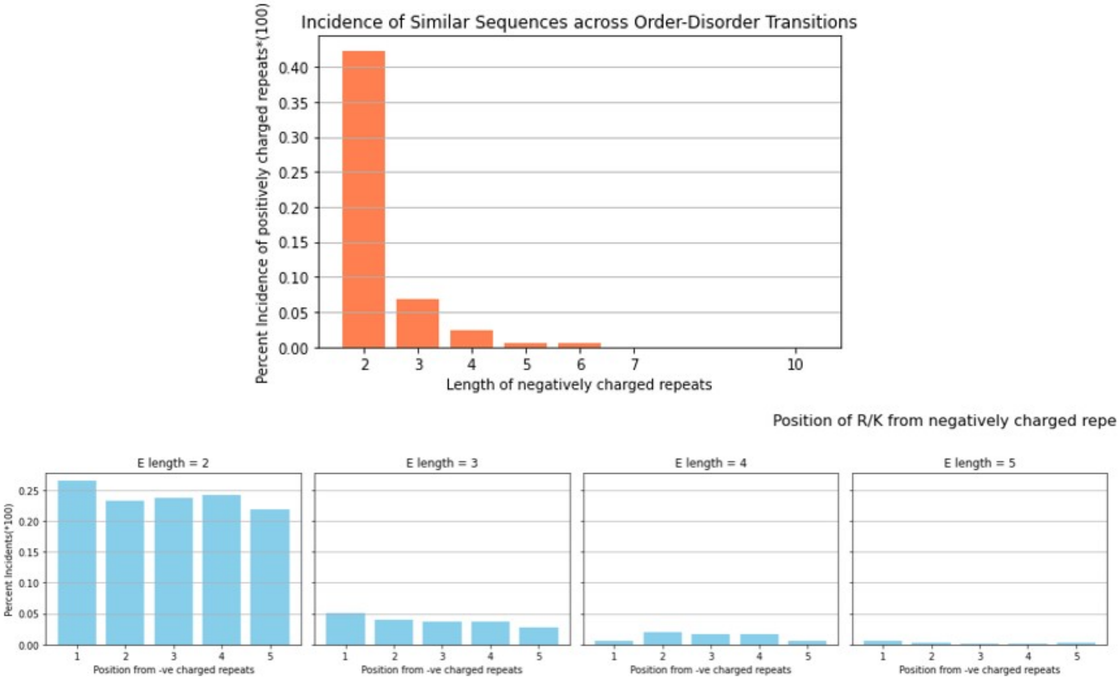


**Supplementary Figure 15: Acidic–basic sequence motifs are prevalent within order–disorder transition regions.** Top, incidence of positively charged residues (R/K) flanking consecutive acidic residue stretches (2–10 E/D residues) within proteins annotated as undergoing order–disorder transitions. Short acidic repeats are the most abundant, with frequency decreasing sharply as repeat length increases. Bottom, positional distribution of R/K residues located 1–5 residues downstream of acidic repeats of varying length (E = 2–5). While short acidic repeats show a broadly uniform distribution of downstream basic residues, longer acidic stretches occur less frequently. These data indicate that sequence architectures resembling the charged–hydrophobic–charged motif of H8 are widely distributed across disordered recognition regions.

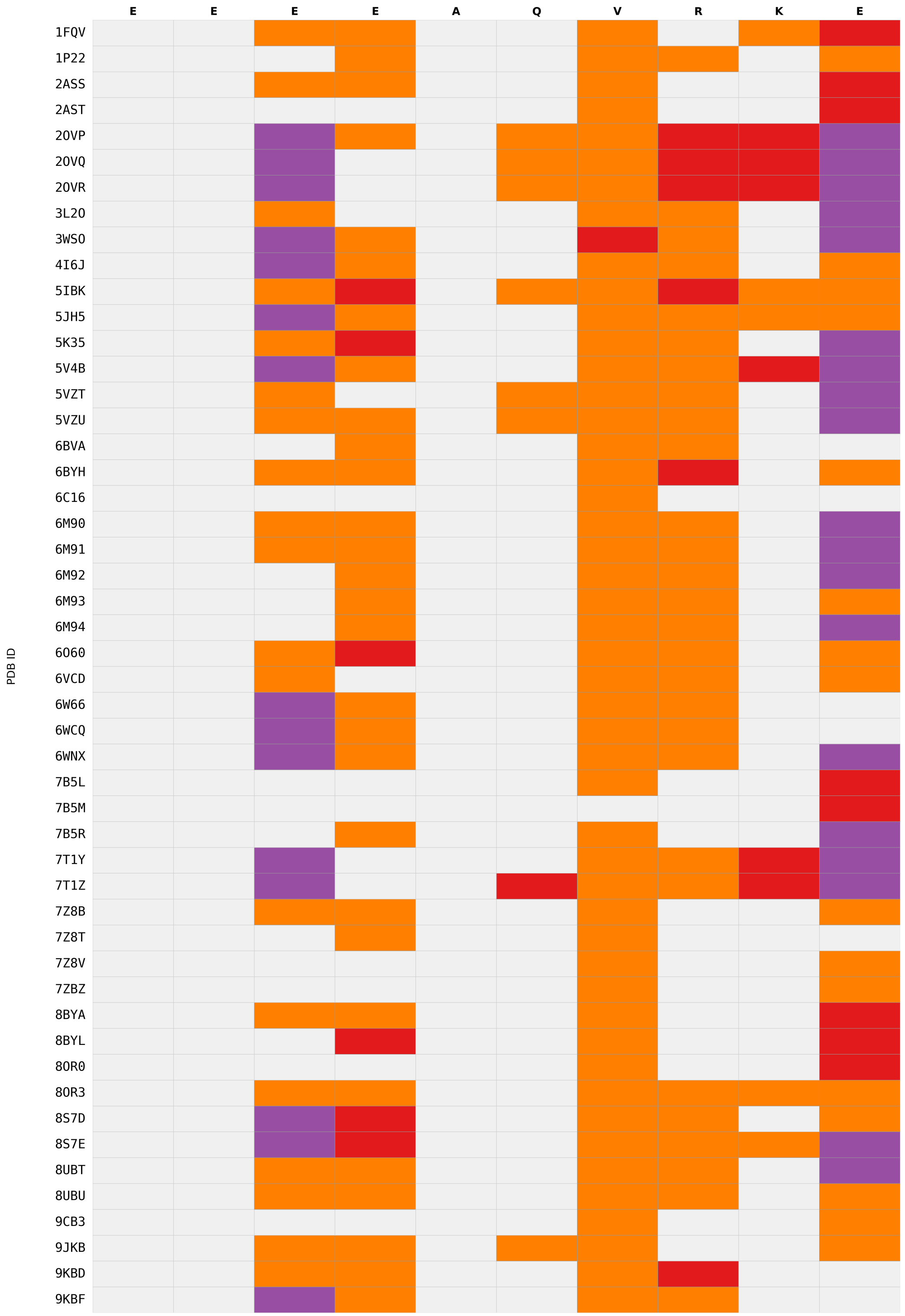


**Supplementary Figure 16: Intermolecular contacts formed by H8 across Skp1-bound structures.** Heatmap summarising residue-specific intermolecular contacts formed by the Skp1 H8 motif (EEEEAQVRKE) across experimentally determined structures. Columns correspond to H8 residue positions and rows correspond to individual PDB structures. Interaction types were assigned from structural coordinates using atom-specific distance criteria: electrostatic contacts between oppositely charged atoms within 4.0 Å, hydrogen bonds between N/O atoms within 3.5 Å excluding salt bridges, and hydrophobic contacts for remaining intermolecular contacts within the distance cutoff. Grey indicates no detected intermolecular contact. The interaction map shows preferential engagement of the C-terminal half of H8, particularly around V153, R154, K155 and E156, consistent with partner recognition involving the stabilised helical face.

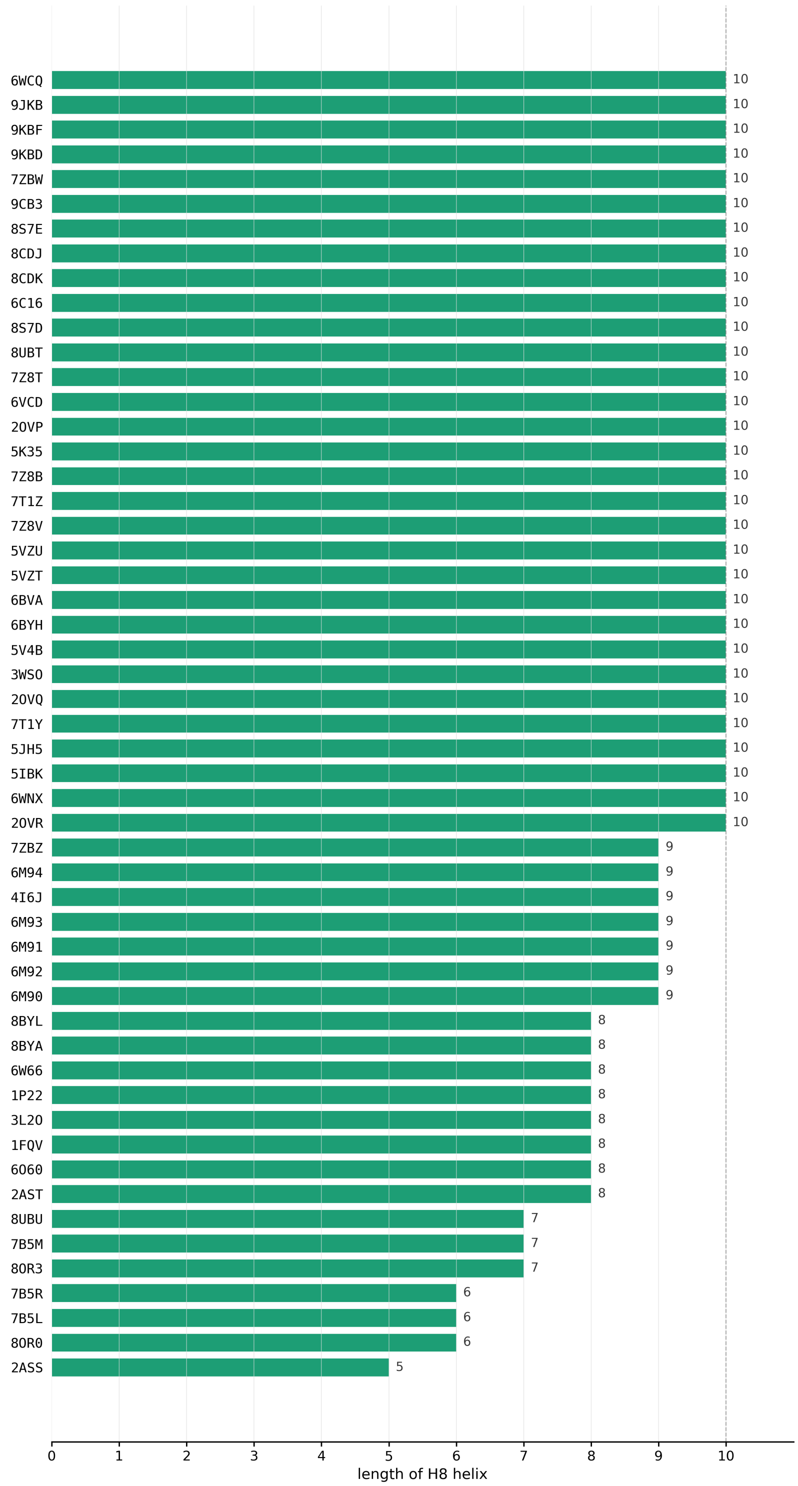


**Supplementary Figure 17: H8 helix length varies across Skp1-containing structures.** Horizontal bar plot showing the number of residues adopting helical secondary structure within the conserved H8 motif (EEEEAQVRKE) across experimentally determined Skp1-containing structures. Secondary-structure assignments were obtained from DSSP v3.0 analysis of mmCIF files, with α-helix, $3_{10}$-helix and π-helix assignments counted as helical. Structures are ordered by increasing H8 helix length. The dashed vertical line indicates the maximum possible helix length of the 10-residue motif, and numerical labels denote the number of helical residues identified in each structure.

**Supplementary tables**

**Supplementary Table 1: Secondary Structure Regions across $Skp1_{helix}$ and $Skp1_{coil}$ conformations.** $Skp1_{helix}$ was prepared as used before by Dantu et al, where the missing regions were modelled (Loops 1 and 2) using PDB ID: 1FQV as a template and $Skp1_{coil}$ was used from PDB ID: 5XYL. Minor changes in the structure are present especially towards the loop regions between helices with major change reflected across the presence and absence of Helix 8.

| [illegible] | [illegible] | [illegible] |
|---|---|---|
| [illegible] | [illegible] | [illegible] |
| [illegible] | [illegible] | [illegible] |
| [illegible] | [illegible] | [illegible] |
| [illegible] | [illegible] | [illegible] |
| [illegible] | [illegible] | [illegible] |
| [illegible] | [illegible] | [illegible] |
| [illegible] | [illegible] | [illegible] |
| [illegible] | [illegible] | [illegible] |
| [illegible] | [illegible] | [illegible] |
| [illegible] | [illegible] | [illegible] |
| [illegible] | [illegible] | [illegible] |
| [illegible] | [illegible] (present) | (absent) |

**Supplementary Table 2: Simulation summary for the $Skp1_{helix}$ ensemble.** Individual trajectory lengths and cumulative simulation time for simulations initiated from the Skp1–Skp2 crystal complex (PDB: 1FQV), in which H8 adopts a helical conformation. Missing loop regions were modelled as described previously (Dantu et al., 2017). The total simulation time was 171.5 µs.

| Replicate | Length(µs) |
|---|---|
| 1 | 1.79 |
| 2 | 1.87 |
| 3 | 1.90 |

| | |
|---|---|
| 4 | 1.90 |
| 5 | 1.88 |
| 6 | 1.89 |
| 7 | 1.89 |
| 8 | 1.88 |
| 9 | 1.82 |
| 10 | 1.89 |
| 11 | 1.88 |
| 12 | 1.88 |
| 13 | 1.87 |
| 14 | 1.89 |
| 15 | 1.00 |
| 16 | 1.00 |
| 17 | 1.88 |
| 18 | 1.91 |
| 19 | 1.87 |
| 20 | 1.89 |
| 21 | 1.89 |
| 22 | 1.91 |
| 23 | 1.90 |
| 24 | 1.91 |
| 25 | 5.41 |
| 26 | 5.36 |
| 27 | 5.43 |
| 28 | 4.66 |
| 29 | 5.47 |
| 30 | 5.44 |
| 31 | 5.47 |
| 32 | 5.48 |
| 33 | 5.40 |
| 34 | 5.43 |
| 35 | 5.42 |
| 36 | 5.32 |
| 37 | 5.42 |
| 38 | 5.41 |
| 39 | 4.64 |
| 40 | 5.36 |
| 41 | 4.55 |
| 42 | 4.59 |

| | |
|---|---|
| 43 | 4.55 |
| 44 | 4.53 |
| 45 | 3.70 |
| 46 | 3.87 |
| 47 | 4.55 |
| 48 | 4.55 |
| 49 | 4.51 |
| 50 | 3.58 |
| **Total length** | **171.51** |

**Supplementary Table 3: Simulation summary for the $Skp1_{coil}$ ensemble.** Individual trajectory lengths and cumulative simulation time for simulations initiated from the ten conformational models of the apo Skp1 solution NMR structure (PDB: 5XYL), in which H8 is predominantly disordered. The total simulation time was 201.1 μs.

| NMR_model | Replicate | Length (μs) |
|---|---|---|
| 1 | 1 | 2.01 |
| 1 | 2 | 1.99 |
| 1 | 3 | 7.55 |
| 1 | 4 | 7.08 |
| 1 | 5 | 7.33 |
| 1 | 6 | 4.91 |
| 1 | 7 | 6.69 |
| 1 | 8 | 3.40 |
| 1 | 9 | 4.58 |
| 1 | 10 | 5.22 |
| 2 | 1 | 2.01 |
| 2 | 2 | 1.99 |
| 2 | 3 | 7.52 |
| 2 | 4 | 6.96 |
| 2 | 5 | 7.37 |
| 2 | 6 | 4.95 |
| 2 | 7 | 6.74 |
| 2 | 8 | 3.45 |
| 2 | 9 | 4.54 |
| 2 | 10 | 5.17 |
| 3 | 1 | 2.02 |
| 3 | 2 | 1.99 |
| 3 | 3 | 7.63 |
| 3 | 4 | 6.98 |
| 3 | 5 | 7.38 |
| 3 | 6 | 4.22 |
| 3 | 7 | 6.72 |
| 3 | 8 | 3.35 |
| 3 | 9 | 4.47 |
| 3 | 10 | 5.13 |
| 4 | 1 | 2.01 |
| 4 | 2 | 1.98 |
| 4 | 3 | 7.61 |

| | | |
|---|---|---|
| 4 | 4 | 6.90 |
| 4 | 5 | 7.37 |
| 4 | 6 | 4.21 |
| 4 | 7 | 6.61 |
| 4 | 8 | 3.33 |
| 4 | 9 | 4.53 |
| 4 | 10 | 5.21 |
| | **Total Length** | **201.10** |

**Supplementary Table 4: Residue-level fractional helicity across H8.** Average fractional helicity for residues 147–156 calculated from DSSP assignments in the $Skp1_{helix}$ and $Skp1_{coil}$ ensembles. Higher values indicate greater residence time in helical conformations and identify residues contributing to transient helix formation within H8.

| Residue | $Skp1_{helix}$ Mean | $Skp1_{coil}$ Mean |
|---|---|---|
| **147** | 0.65 | 0.2 |
| **148** | 0.83 | 0.24 |
| **149** | 0.93 | 0.33 |
| **150** | 0.93 | 0.55 |
| **151** | 0.92 | 0.47 |
| **152** | 0.88 | 0.31 |
| **153** | 0.75 | 0.18 |
| **154** | 0.51 | 0.09 |
| **155** | 0.34 | 0.16 |
| **156** | 0.15 | 0.17 |

**Supplementary Table 5: Mean transition times between Markov state modelling (MSM) macrostates.** Mean first-passage times between H8 macrostates identified from MSM. Macrostates M0 and M4 correspond to the dominant coil basin, M3 corresponds to the rare fully helical state, and M1, M2 and M5 represent intermediate partially helical states connecting the coil and helical basins. Populations of the initial and final states are shown alongside transition times in microseconds.

| **From state** | **To state** | **Population$_i$** | **Population$_j$** | **Mean transition time (in µs)** |
|---|---|---|---|---|
| **M0** | M1 | 0.44 | 0.00 | 788.77 |
| **M0** | M2 | 0.44 | 0.17 | 9.35 |
| **M0** | M4 | 0.44 | 0.37 | 2.83 |
| **M0** | M5 | 0.44 | 0.02 | 161.33 |
| **M1** | M0 | 0.00 | 0.44 | 4.35 |
| **M1** | M2 | 0.00 | 0.17 | 5.15 |
| **M1** | M4 | 0.00 | 0.37 | 4.78 |
| **M1** | M5 | 0.00 | 0.02 | 21.25 |
| **M2** | M0 | 0.17 | 0.44 | 3.37 |
| **M2** | M4 | 0.17 | 0.37 | 3.83 |
| **M2** | M5 | 0.17 | 0.02 | 26.39 |
| **M3** | M0 | 0.00 | 0.44 | 5.12 |
| **M3** | M1 | 0.00 | 0.00 | 6.51 |
| **M3** | M2 | 0.00 | 0.17 | 7.67 |

| | | | | |
|---|---|---|---|---|
| **M3** | M4 | 0.00 | 0.37 | 5.20 |
| **M3** | M5 | 0.00 | 0.02 | 20.22 |
| **M4** | M0 | 0.37 | 0.44 | 2.42 |
| **M4** | M2 | 0.37 | 0.17 | 7.57 |
| **M4** | M5 | 0.37 | 0.02 | 91.00 |
| **M5** | M0 | 0.02 | 0.44 | 3.65 |
| **M5** | M1 | 0.02 | 0.00 | 66.55 |
| **M5** | M2 | 0.02 | 0.17 | 4.36 |
| **M5** | M3 | 0.02 | 0.00 | 64.27 |
| **M5** | M4 | 0.02 | 0.37 | 5.33 |

**Supplementary Table 6: E-CPMG exchange kinetics for H8 residues at 272 K.** Residue-specific exchange rate constants (Kex) obtained from E-CPMG relaxation-dispersion analysis at 272 K for H8 residues showing detectable conformational exchange. Reported values are given as Kex × $10^3$ $s^{-1}$ with associated fitting uncertainties.

| **Residue** | **$K_{ex}$ ($10^3$ $s^{-1}$)** |
|---|---|
| **E150** | 27.66 ±10.67 |
| **A151** | 38.54 ± 9.91 |
| **Q152** | 33.31 ± 3.83 |
| **V153** | 14.58 ± 2.14 |
| **R154** | 26.24 ± 20.19 |
| **K155** | 5.82 ± 1.20 |

**Supplementary Table 7: Experimentally determined structures of Skp1 in complex with F-box proteins (FBPs).** List of Skp1-containing structures deposited in the Protein Data Bank, including complexes with diverse FBPs and the apo solution NMR structure of Skp1 (PDB: 5XYL). These structures were used to analyse H8 conformational variability and intermolecular interactions associated with F-box protein recognition.

| PDB ID | Bound Protein |
|---|---|
| 1FS1; 2AST; 7Z8V; 1FQV; 1FS2; 2ASS; 2AST; 7Z8T; 1LDK; 7ZBZ; 8OR0; 8CDK; 8BYA; 8CDJ; 7ZBW; 8BYL; 7B5L; 7B5R; 8OR4; 7B5M; 8OR3 | **Skp2** |
| 6M90; 6M91; 6M92; 6M93; 6M94; 1P22; 6TTU | **β-TRCP1** |
| 2OVP; 2OVR; 2OVQ; 5IBK; 7T1Y; 7T1Z; 5V4B | **FBXW7** |
| 6WNX | **FBXW11** |
| 6O60 | **FBXL2** |
| 5JH5; 6BVA | **FBXL10** |
| 3WSO | **FBXO44** |
| 6BYH; 6C16 | **FBXL11** |
| 4I6J | **FBXL3** |
| 5VZT; 5VZU | **FBXO31** |
| 3L2O | **FBXO4** |
| 7Z8B | **FBXW8** |
| 6VCD | **FBXL5** |
| 8S7D; 8S7E; 8UA6 | **FBXO22** |
| 8UBT; 8UBU; 6WCQ; 6W66 | **FBXL17** |
| 9CB3 | **Cyclin F** |
| 9KBD; 9KBF | **FBXO3** |
| 9JKB | **FBXO4** |
| 5XYL (APO) | **Skp1** |
